# A translational repression reporter assay for the analysis of RNA-binding protein consensus sites

**DOI:** 10.1101/2022.09.05.506580

**Authors:** Jessica Nowacki, Mateo Malenica, Stefan Schmeing, Damian Schiller, Benjamin Buchmuller, Peter ‘t Hart

## Abstract

RNA-binding proteins are essential regulators of RNA processing and function. Translational repression assays can be used to study how they interact with specific RNA sequences by insertion of such a consensus sequence into the 5’ untranslated region of a reporter mRNA and measuring reporter protein translation. The straightforward set-up of these translational repression assays avoids the need for the isolation of the protein or the RNA providing speed, robustness, and a low-cost method. Here we report the optimization of the assay to function with linear RNA sequences instead of the previously reported hairpin type sequences to allow the study of a wider variety of RNA-binding proteins. Multiplication of a consensus sequence strongly improves the signal allowing analysis by both fluorescence intensity measurements and fluorescence activated cell sorting.

## INTRODUCTION

RNA-binding proteins (RBPs) are essential in the processing and function of all RNA types and therefore play a prominent role in physiology and disease.(1–3) Studying how RBPs interact with specific RNA sequences is relevant to learn more about their function and selectivity and to identify novel therapeutic targets.(4, 5) Experiments using purified proteins and RNA can lead to novel insights but can be affected by the lack of the components of the intracellular environment. Furthermore, obtaining both protein and RNA in good purity and sufficient quantities can be challenging and laborious. These challenges can be overcome by performing protein-RNA interaction assays in bacterial cells and various assays have been reported to do so including the bacterial three-hybrid assay, antitermination based assays, and assays based on translational repression.(6–10) Although the bacterial three-hybrid assay has been shown to be effective at coupling a protein-RNA interaction to a read-out, it requires three components to be constructed and has been reported to only be effective for high-affinity interactions.(7) The antitermination-based assays are hard to adapt to new protein-RNA interactions due to conformational restrictions and also require very high affinities to produce a read-out.(11, 12) The systems based on translational repression require only two designed components and have been demonstrated to work in bacterial, yeast, and mammalian cells.(13–17) In contrast to the other two systems, the RBP is conformationally free to interact with the target RNA rather than being part of a larger complex making the system more flexible and straightforward to design. Various read-outs have been tested in such repression assays including survival-based selectors, or read-outs based on colorimetric (i.e.: LacZ), luminescent (i.e.: luciferase), or fluorescent reporter proteins (i.e.: GFP). A major advantage of the translational repression assay is that the repression levels appear to correlate with binding affinity allowing the study of protein-RNA interactions without purification of the individual components.(10, 17)

The translational repression system relies on an mRNA encoding for a reporter gene which is modified with an RBP binding site (see Fig. 1A) by inserting it in the 5’ UTR upstream of the ribosomal binding site.(14, 15, 18) Alternatively, insertion of the RBP recognized sequence after the start codon and in frame with the reporter protein has been reported as well.(16, 17) Binding of an RBP to the RBP binding site leads to a blockade of reporter protein translation which can be used as a measure for the protein-RNA interaction (see Fig. 1B). By fusing the RBP to a fluorescent protein compatible with the reporter protein, the RBP expression levels can be monitored simultaneously. In previously described versions of the system, the RBP binding site has always been a stable hairpin RNA to facilitate a high affinity interaction (e.g. MS2 with the MS2 hairpin).(14) However, in our studies we are interested in alternative splicing factors that interact with unstructured linear RNA sequences. Until now, only the core splicing component U1A has been tested in this assay, which has a very high affinity for the U1 snRNA hairpin. We therefore sought to adapt the system, so it accepts such interactions and allow straightforward modification and interaction analysis.

**Figure 1.**
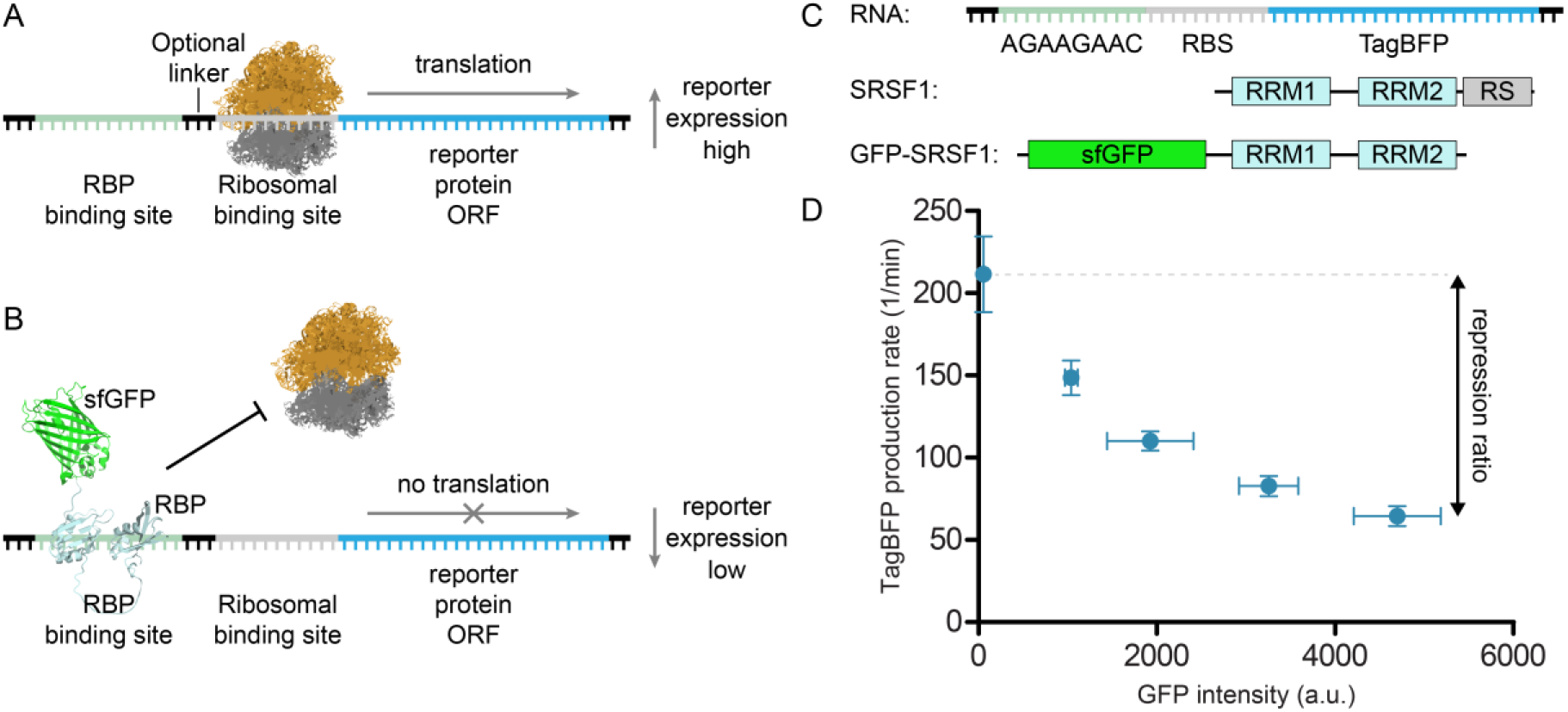
Translational repression assay. A) In the absence of the RBP, translation of the reporter protein can take place. B) In the presences of the RBP-sfGFP fusion construct, translation is blocked. C) Schematic representation of the modified TagBFP mRNA reporter used as starting point for assay development, schematic representation of the SRSF1 domain structure, and schematic representation of the used SRSF1 protein construct GFP-SRSF1. D) Repression of TagBFP observed upon increasing expression of GFP-SRSF1 The repression ratio is calculated by dividing the highest with the lowest TagBFP production rates.

Here we report a variation of the translational repression system that allows the study of the interaction between an RBP and a linear RNA sequence. The RNA sequence is inserted in front of the ribosomal binding site of a blue fluorescent protein (TagBFP) reporter so that RBP binding interferes with TagBFP translation. We found that TagBFP repression can be increased by multiplication of the RNA sequence and varying the distance between the insert and ribosomal binding site. We demonstrate the use of the assay for two different splicing factors (SRSF1 and hnRNP A2/B1) and used both a plate reader-based read-out as well as flow cytometry. Binding of the RBPs to the used sequences was further verified using fluorescence polarization assays and increased repression correlated with binding affinity. Our findings demonstrate that the assay robustly reports on binding affinity, but for a strong signal-to-noise ratio it is important to make sure no secondary structures are formed in the 5’-UTR of the reporter in close proximity to the ribosomal binding site.

## MATERIAL AND METHODS

### Protein expression and purification

MBP-tagged hnRNPA2B1 (aa1-251) was expressed at the Protein Chemistry Facility (PCF) based in the MPI Dortmund. HnRNPA2B1 (1-251) was sub-cloned into pOPIN-His-MBP multihost expression vectors. The MBP fusion was chosen to increase expression yield and solubility.(19) MBP-hnRNPA2B1 was expressed in E.coli BL21 CodonPlus (DE3) RIPL. Bacteria with the respective plasmid were cultured in Terrific Broth with 0.01% lactose, 2 mM MgSO_4_, 100 µg/ml ampicillin and 50 µg/ml chloramphenicol. Expression of protein was auto-induced, with incubation of the starter-culture (starting OD of ∼0.05) at 37 °C for 4 h, followed by an overnight incubation (20-24h) at 25 °C. Bacteria were harvested by centrifugation and lysed (50 mM HEPES, 300 mM NaCl, 20 mM Imidazole, 1 mM TCEP, pH 8) using a Celldisrupter TS 0.75 (Constant Systems) at 1350 bar.

Protein purification of MBP-tagged hnRNPA2B1 was also performed at PCF. Protein purification was performed on HisTrap FF crude 5 ml Ni-based column using ÄKTA Xpress System (Cytiva, former GE Healthcare). For that a wash buffer (50 mM HEPES, 300 mM NaCl, 30 mM Imidazole, 1 mM TCEP at pH 8) and elution buffer (50 mM HEPES, 300 mM NaCl, 500 mM Imidazole, 1 mM TCEP at pH 8) were used. The protein was then further purified using size exclusion chromatography (HiLoad 26/60 Superdex 75 prep grade column) in storage buffer (50 mM HEPES, 100 mM NaCl, 1 mM TCEP at pH 8.0) at 4 °C. Fractions were collected and concentrated with a 50 kDa molecular mass cutoff Amicon.

SRSF1 RRM1+2 (aa1-195) was sub-cloned into pOPIN-His multihost expression vectors by SLIC and expressed in E.coli BL21 (DE3) and purified using a protocol adapted from Clery et al.(19, 20) Bacteria with the respective plasmid were cultured in LB medium with 100 µg/ml ampicillin. Protein expression was induced at OD_600_=0.6 with 1 mM isopropyl β-D-thiogalactoside (IPTG) overnight at 18°C. Cells were harvested by centrifugation and lysed (50 mM Na_2_HPO_4_, 300 mM KCl, 50 mM L-Arg, 50 mM L-Glu, 1.5 mM MgCl_2_, 1 mM PMSF at pH 8) using a microfluidizer. Protein purification was performed on HisTrap HP 5 ml Ni-based column (Cytiva, former GE Healthcare) using ÄKTA Explorer System (Cytiva, former GE Healthcare). The protein was dialyzed overnight at 4°C into wash buffer (50 mM Na_2_HPO_4_, 300 mM KCl, 50 mM L-Arg, 50 mM L-Glu, 1.5 mM MgCl_2_, 40 mM Imidazole at pH 8), and a second Ni-based column purification was performed. The protein was again dialyzed at 4°C into wash buffer and treated with His-tagged 3C protease. After overnight cleavage the protein was loaded onto Ni-based column for reversed purification. The protein was finally dialyzed at 4 °C in the storage buffer (20 mM NaH_2_PO_4_, 150 mM KCl, 50 mM L-Arg, 50 mM L-Glu,m 1.5 mM MgCl_2_, 0.2 mM EDTA, 1 mM TCEP at pH 7) and concentrated with a 3 kDa molecular mass cutoff Amicon.

### Translational Repression Assay Procedure

A 1 mL preculture of E.coli Top10F’ cotransformed with assay plasmids in LB with corresponding antibiotics (Kanamycin 50 µg/mL, Chloramphenicol 34 µg/mL) in a 96-deep well plate was prepared. The plate was sealed with a semipermeable sticky lid and the cultures were grown overnight at 37 °C in a shaking incubator (160 rpm). The next day the cultures were diluted 1:19 in M9 minimal medium with respective antibiotics in a total volume of 190 µL in a black, 96-well plate with clear bottom. Optical density at 600 nm (OD600) was monitored until a value of ∼0.2 was reached. The assay was induced with IPTG and arabinose to final concentrations of 1 mM and 0, 0.125, 0.25, 0.5 or 1 %. Fluorescence of TagBFP (402 nm/457 nm) and sfGFP (485 nm/510 nm) in addition to OD_600_ were measured every 20 minutes over a time course of ∼ 7 h at 30 °C with a TECAN Spark plate reader. Data evaluation was performed according to the report by Katz et al.(21)

### Flow cytometry analysis

A 1 mL preculture of E.coli Top10F’ cotransformed with assay plasmids in LB medium with corresponding antibiotics (Kanamycin 50 µg/mL, Chloramphenicol 34 µg/mL) in a 96-deep well plate was prepared. The plate was sealed with a semipermeable sticky lid and the cultures were grown overnight at 37 °C in a shaking incubator (160 rpm). The next day the cultures were diluted 1:19 in LB medium with respective antibiotics in a total volume of 1 mL in a 96-deep-well plate. OD_600_ was monitored until a value of ∼0.2 was reached. The assay was induced with 1mM IPTG and 0.125 % arabinose for 4 h and 30 minutes at 30 °C. The plate was centrifuged at 4500 g for 5 minutes in a tabletop centrifuge. Medium was removed and cells were washed twice with 500 µL PBS. Finally, cells were centrifuged and resuspended in 800 µL PBS and placed on ice until analysis.

Samples were analyzed on a SH800SFP Cell Sorter (Sony Biotechnology, Weybridge, U. K.) using a 70 μm microfluidic chip (Sony Biotechnology), and TagBFP and sfGFP fluorescence intensities recorded and compensated using the respective single-color controls (see supplemental figure 35).”

### Fluorescence polarization Assay

The RNA oligonucleotides were purchased by Integrated DNA Technologies or SigmaAldrich. Oligonucleotides were dissolved in nuclease-free water to 100 µM according to the manufacturers’ instructions and kept at -20 °C until use. The assay was performed in protein storage buffer with 0.01% Triton, in black, 384 well-plates (Corning), with a total volume of 20 μL per well. The analyzed, 6-FAM labeled RNAs were tested at a final concentration of 1 nM, and the appropriate unlabeled protein was titrated as two-fold dilution series. The plates were analyzed subsequently in a plate reader (Tecan Spark) using (ex/em) 490/520 nm for 6-FAM. The assay was performed in two duplicates on two independent days (quadruplicates).

## RESULTS

### Development of the assay for SRSF1

To test the use of linear sequences in the translational repression system we initially used SRSF1 as our RBP of choice. SRSF1 is a splicing factor that interacts with exonic splicing enhancer (ESE) sequences to influence splicing events.(22, 23) Two RNA-recognition motif (RRM) domains drive its RNA binding and are followed by a *C*-terminal unstructured arginine-serine rich (RS) domain (see Fig. 1C).(20, 22) To prepare an SRSF1 construct for use in the translation repression assay we fused sfGFP to the SRSF1 *N*-terminus (see Fig. 1C) and removed the RS domain as this was previously shown not to participate in RNA binding (GFP-SRSF1).(24) As the RBP binding site in the TagBFP reporter we chose a consensus sequence (AGAAGAAC) previously identified to be recognized by SRSF1 via SELEX experiments.(25) In contrast to the method described by Katz *et al*. we introduced the RBP binding site in front of the RBS rather than after the start codon. By doing so we avoided the need for a design that stays in frame with the TagBFP ORF. The sequence was inserted directly in front of the ribosomal binding site (see Fig. 1C) and both sfGFP and TagBFP intensity were measured under increasing concentrations of the inducing agent (i.e. arabinose) for GFP-SRSF1 while the RNA was constantly induced with IPTG (1 mM) (see Fig. 1D). To measure repression, we used the method described by Katz *et al*. which determines the fluorescent protein production rate rather than steady state levels to avoid saturation of the system.(17, 21) The TagBFP production rates are plotted against the sfGFP expression levels. TagBFP production rate is represented by the ratio of TagBFP levels divided by the integral of cell density in a time interval within the linear growth phase. sfGFP expression levels were normalized to cell density and averaged by the number of time points within the chosen time interval. To facilitate straightforward comparison of different RNA-RBP pairs we calculated a repression ratio by dividing the baseline TagBFP production rate (non-induced sfGFP-RBP) by the TagBFP production rate at the highest expression level (at 1% arabinose) of the sfGFP-RBP fusion (see Fig. 1D). Such repression ratios have been used previously and allow straightforward comparison of different RNA/RBP pairs.(14, 18, 26)

In the first reporter construct S1 (see Fig. 1C/D and table 1) we introduced the SRFS1 RNA consensus sequence directly upstream of the RBS and we measured a repression ratio of 3.3 ± 0.3. As a control we fused sfGFP to the RRM domains 3 and 4 of the RNA-binding protein PTBP1 (GFP-PTBP1) which recognizes polypyrimidine sequences rather than the purine rich S1 sequence. Expression of this protein led to a 1.5 ± 0.2 fold repression (see supplemental table 1 and supplemental figure 1) which likely represents increased competition for protein synthesis resources. Although repression was stronger in the presence of the SRSF1 construct, a bigger dynamic range would allow a clearer discrimination. The Saito group demonstrated that duplication of a hairpin shaped RBP binding site improved translational repression of a reporter gene.(27) Furthermore, a recent report describing oligonucleotides with potent SRSF1 binding indicated that repeating a consensus sequence two or more times significantly improved binding affinity.(28) To explore whether this phenomenon also improved translational repression in our system we multiplied the RNA sequence two or three times and observed a significant increase in repression ratio of 9.0 (construct S2) and 15.1 (construct S3) as shown in table 1. The increase amounted to more than the sum of the repression ratio observed for a single insert possibly indicating an avidity effect.

**Table 1.**
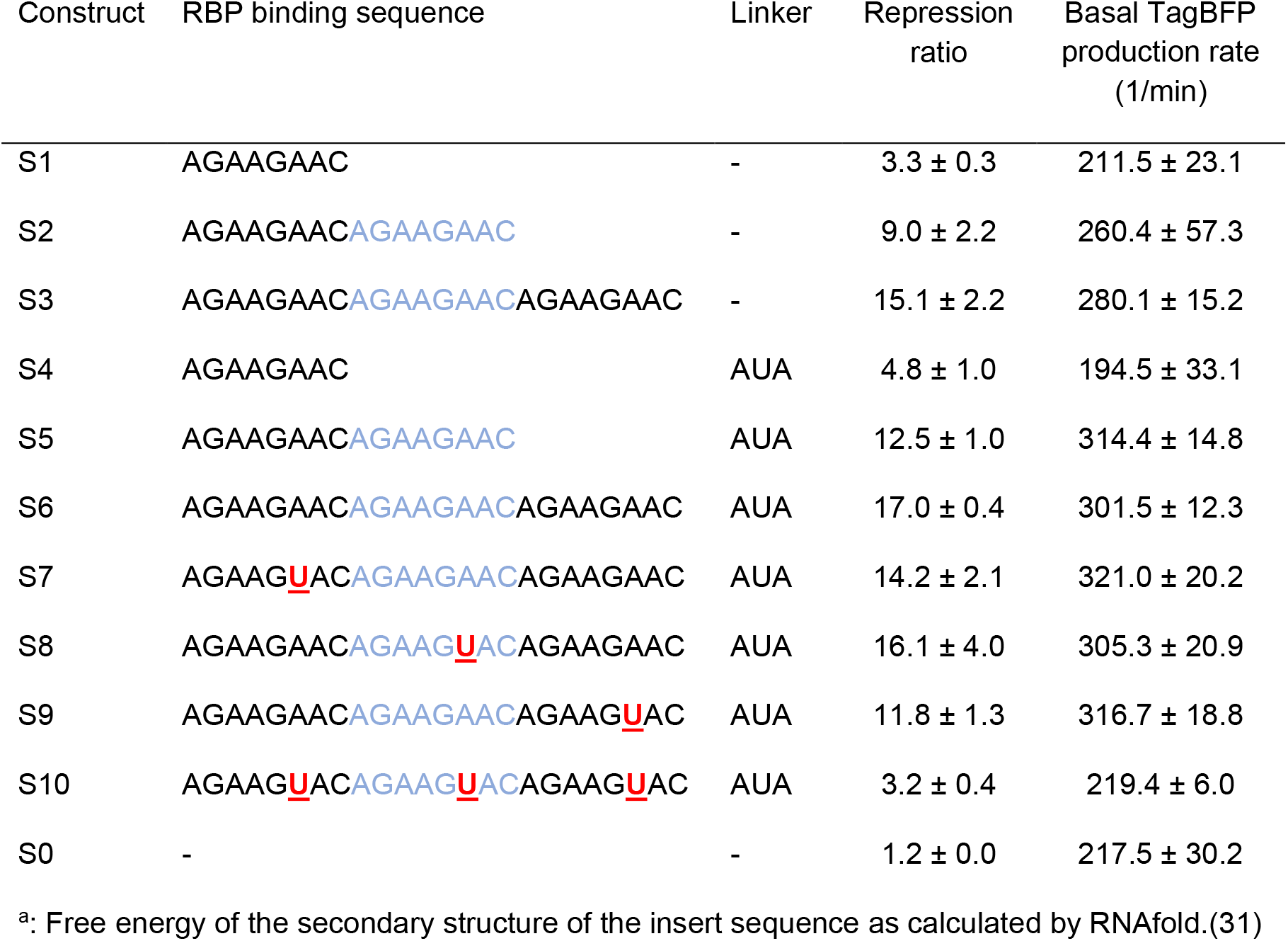
Different reporter constructs used for translational repression assays with SRSF1.

Previous descriptions of translational repression systems with similar designs reported that the distance between the RBP binding site and the RBS can influence the repression.(17) We therefore introduced a short three nucleotide linker to see whether the repression effect improves. Addition of the AUA linker indeed improved the repression for all three different constructs (S4-S6) albeit to a small degree (see Table 1 and supplemental figure 2). Since the S6 sequence displayed the highest repression ratio we tested it in the presence of an SRSF1 mutant (GFP-SRSF1_mut_) we designed to have a strongly reduced affinity for RNA by introducing previously reported mutations in both RRMs, i.e.: F56D and F58D in RRM1, and Q135A and K138A in RRM2.(20, 29) For this combination we observed a repression ratio of 1.6 ± 0.2 (see supplemental table 1 and supplemental figure 2) which is similar to the repression ratio observed for GFP-PTBP1 with this reporter (1.7 ± 0.1, see supplemental table 1) indicating that the repression is driven by RBP binding. As another control we tested repression of the unmodified reporter plasmid S0 in the presence of GFP-SRSF1 and measured a repression ratio of 1.2 ± 0.0.

Next, to gain insight into how multiplication of the binding sequences affected the interaction we introduced a point mutation in the S6 sequence. We prepared variants where the mutation occurred at either the first, second, or third repeat (S7-S9, see Table 1 and supplemental figure 3) while the other two repeats maintained the original sequence. These constructs indicated that if the mutation was in the copies more distant from the ribosomal binding site that the effect was minimal. However, when the mutation was right next to the ribosomal binding site the repression ratio dropped to a value similar to that of S5. When the mutation was inserted in all repeats (construct S10) the repression was significantly decreased and similar to S1. However, the S10 sequence did have some propensity to form a hairpin like structure which could influence this repression value (see supplemental figure 10). All RNA inserts were also tested for translational repression in the presence of GFP-PTBP1 as described above and no significant repression was observed (see supplemental table 1 and supplemental figures 1-3).

### Development of the assay for hnRNP A2/B1

To explore the general applicability of using linear RNA sequences as RBP recognition sites in the translational repression assay we also explored the splicing factor hnRNP A2/B1. Similar to the SRSF1 construct, we prepared an *N*-terminally sfGFP fused construct containing the residues 1-251 (GFP-A2/B1) which are similar to a truncation that was previously described to have high binding affinity for various consensus sequences as determined by isothermal titration calorimetry (see Fig. 3A).(30) We chose to use the RNA sequence AAGGACUAGC which was previously reported to have an affinity of 26.5 nM.(30) The sequence was introduced in the same position in the TagBFP reporter mRNA as single, double, and triple repeats as we did for the SRSF1 sequence and repression ratios were measured (H1-H3, see Table 2 and supplemental figure 4). Surprisingly, multiplication of the recognition sequence did not improve the repression ratio as was observed for the SRSF1 system. Adding the AUA linker (H4-H6, see Table 2 and supplemental figure 5) as done for the S-constructs did not improve the repression ratio and still no trend was observable. It is possible that direct connection of the consensus sequence does not provide enough spacing for two sfGFP-A2/B1 molecules to bind so we inserted a GGG spacer in between two replicated sequences (H7). Construct H7 indeed demonstrated an improved repression ratio over construct H2 and adding the AUA linker (H8) improved it even further (see Table 2 and supplemental figure 6). However, a similar improvement was not observed for the introduction of a spacer in the triplicate sequence (compare H6 and H9 in Table 2 and supplemental figures 5 and 6). Introducing a mutation (sequence H10 and H11, see Table 2 and supplemental figure 7) that was previously reported to reduce the affinity approximately 8-fold, also indeed reduced the repression ratio (compared to H8). Again, as a control we measured the repression of the unmodified TagBFP reporter plasmid in the presence of GFP-A2/B1 (entry H0, see Table 2) and measured a repression ratio of 1.3 ± 0.2 indicating that the reporters do respond to hnRNP A2/B1 binding. Furthermore, all constructs were evaluated in the presence of the non-binding GFP-PTBP1 resulting in repression ratios varying from 1.1 – 1.8 further indicating the reporters do respond to hnRNP A2/B1 binding (see supplemental figures 4-7).

**Figure 2:**
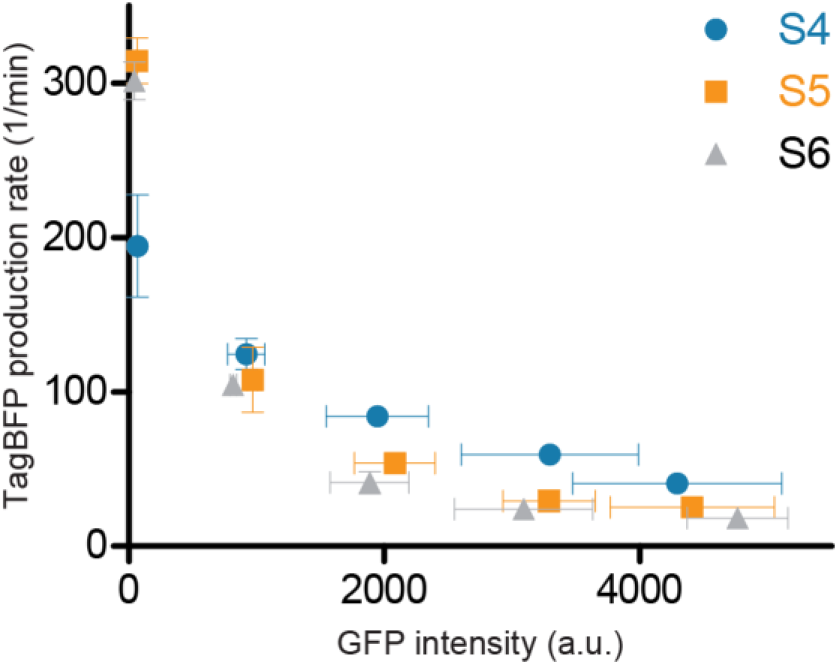
Repression curves of constructs S4-S6 in the presence of increasing concentrations of GFP-SRSF1.

**Figure 3:**
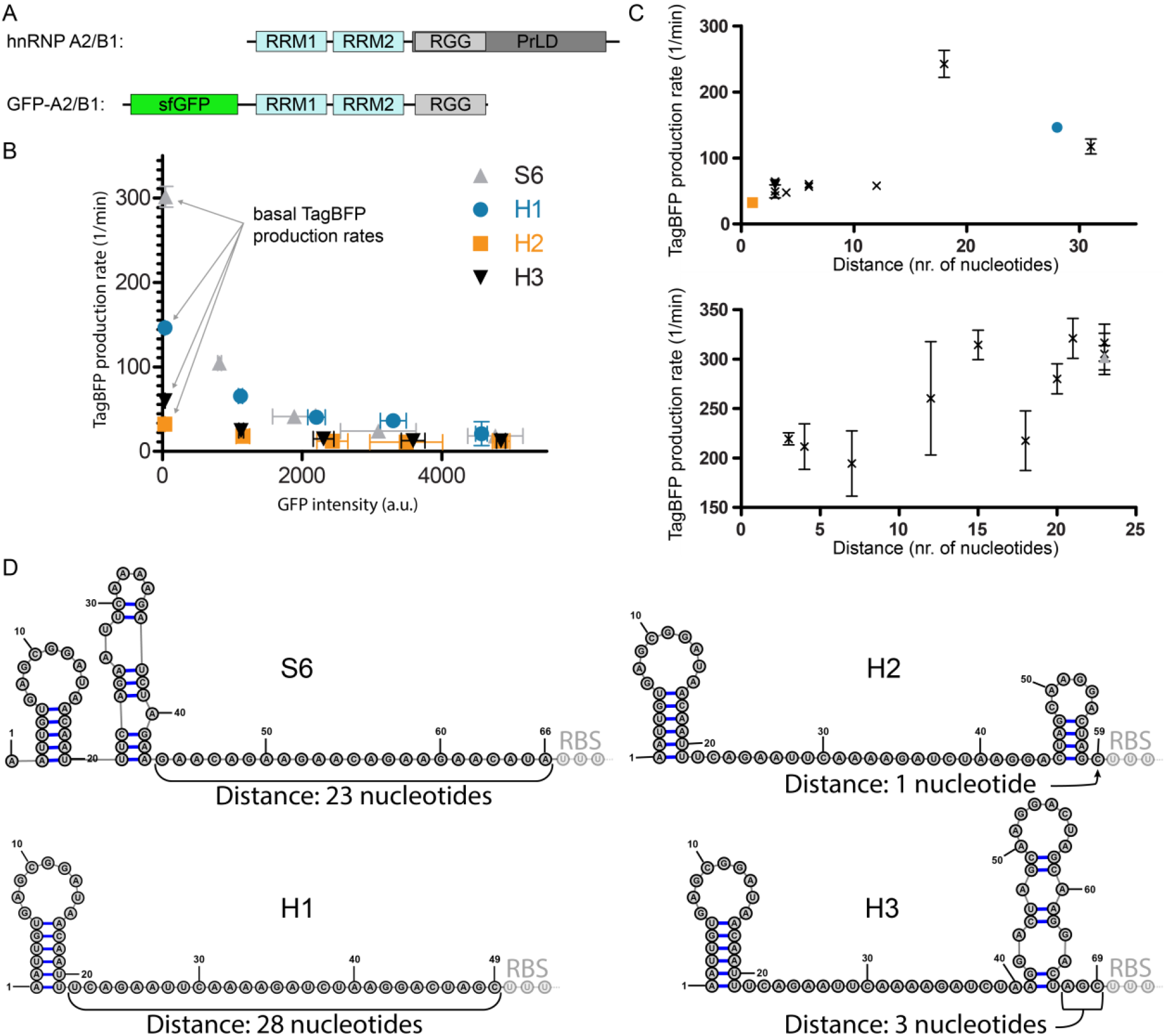
A) Schematic representation of hnRNPA2/B1 and the GFP-A2/B1 protein construct. B) Translational repression assay results for RNA reporters H1-H3 in the presence of GFP-A2/B1 in comparison with RNA reporter S6 in the presence of GFP-SRSF1 C) Correlation between TagBFP production rate and the distance between a secondary structure and the ribosomal binding site for GFP-A2/B1 (top) and GFP-SRSF1 (bottom). D) Predicted secondary structures of the inserted RBP binding sequences for 5’-UTRs of the reporters S6 and H1-H3 using RNAfold. For S6 the first three nucleotides of the RBS sequence are indicated as well as the distance between the RBS and the closest predicted secondary structure.

**Table 2:**
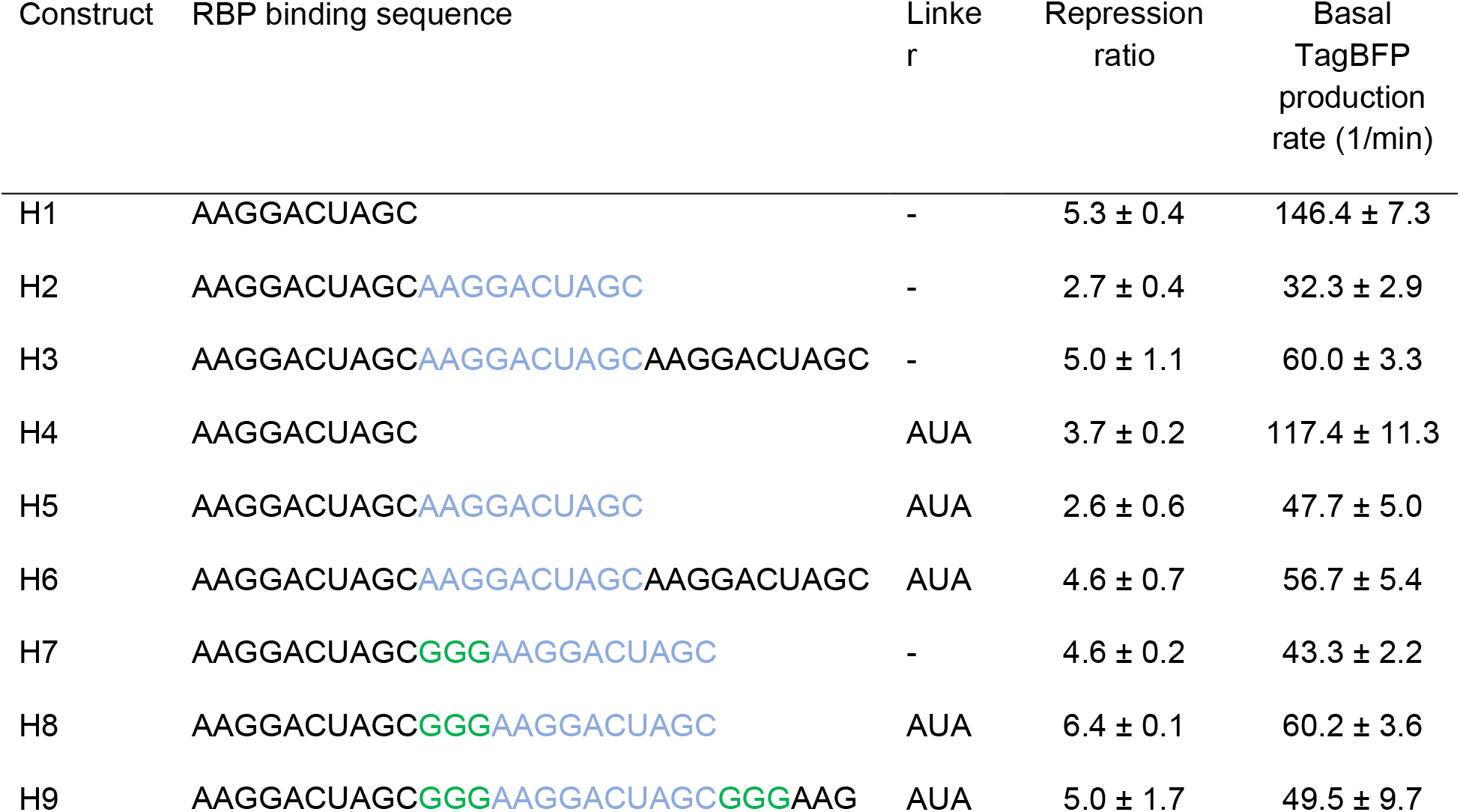
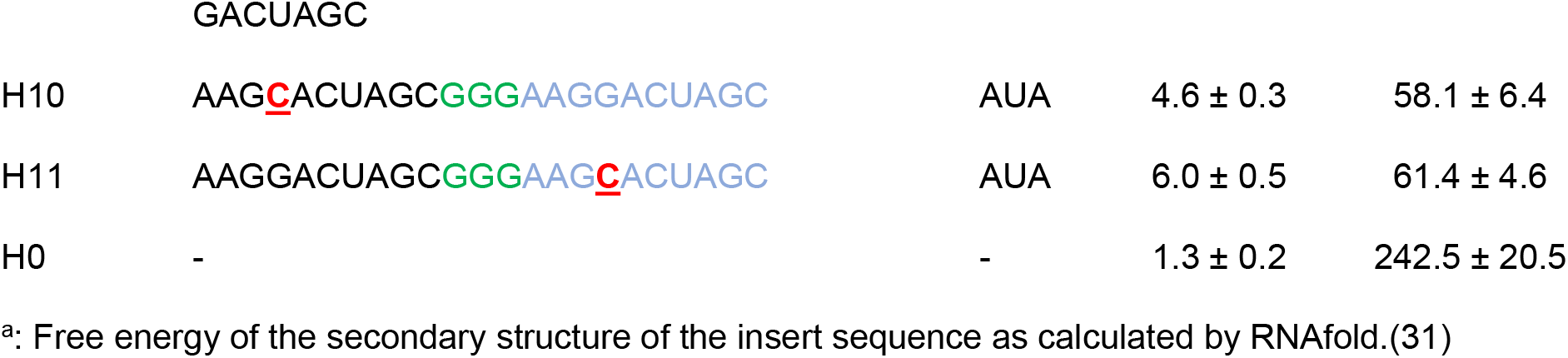
Different reporter constructs used for translational repression assays with hnRNP A2/B1.

### Analysis of the effect of secondary structures in the 5’-UTR of the reporter mRNA

We observed that for several of the H-reporters the basal TagBFP production rate was significantly lower than for most of the S-reporters (see Table 1 and 2, and Fig. 3B). Both the S0 and H0 experiments showed that the basal TagBFP production rate was above 200 1/min, and while most S-reporters were above this level, most H-reporters were significantly below it. The lowered basal TagBFP production rate could potentially be caused by secondary structure formation, and we therefore used the RNAfold algorithm to predicted whether any secondary structures formed in the 5’-UTR of the reporter RNA up until the ribosomal binding site.(31) In all 5’-UTRs there was at least one hairpin but several contained more and in different locations along the sequence (see Fig.3D and supplemental figures 8-15). When the distance between any formed secondary structure and the RBS was plotted against the basal TagBFP production rate a trend became evident that showed that a longer spacing is favored (see Fig. 3C). Although the differences were not so big for the S-reporters, the H-reporter series shows there is a maximum around 18 nucleotides. These results indicate that a reporter should be designed so that it provides a high basal production and therefore sufficient dynamic range to analyze changes in RNA-binding after mutation.

### Correlation of translational repression with binding affinity

To investigate whether increased repression correlated with binding affinity we performed fluorescence polarization (FP) experiments with fluorescein amine (6-FAM) labeled RNA sequences and recombinantly expressed SRSF1 and hnRNP A2/B1 proteins spanning the same residues as used in the translational repression assay but without the *N*-terminal GFP fusion. Instead, hnRNP A2/B1 was modified with an MBP-tag to improve yields during expression and purification while SRSF1 was untagged (see supplemental table 4). For SRSF1 we tested the RNA sequences equivalent to S1, S2, and S3 and for hnRNP A2/B1 the sequences used for H1, H2, and H3 (see Table 3 and supplemental figures 18 and 19). For SRSF1 the results follow the logic of the repression experiments where a single repeat has a low affinity, but a double and triple replication leads to a significantly improved affinity presumably due to a cooperativity effect. The hill-slopes of the measured curves indicate a binding ratio where more than one copy of SRSF1 binds to the RNA sequences. The very low affinity for a single repeat is likely due to the RNA sequence being too short to be effectively bound by the two SRSF1 RRMs. In the repression assay this might be compensated by the flanking sequences of the reporter mRNA. The FP experiments for hnRNP A2/B1 also correlate with the repression assay where H1 and H3 show good binding affinity but a reduction for H2 is observed. The results are somewhat unexpected but are likely caused by secondary structure formation and we therefore also predicted the structures of these isolated RNA sequences using RNAfold (see supplemental figures 16 and 17). Indeed, both FAM-H1 and FAM-H3 possess the AAGGACU sequence in non-base paired form which is recognized by the two RRMs according to the reported crystal structure.(30) In contrast, the binding sequence is embedded into the stem of FAM-H2 probably leading to a balance between the energy required to disrupt this and the binding energy for the linear sequence explaining its reduced affinity.

**Table 3.** Affinities of SRSF1 and hnRNP A2/B1 for representative RNA sequences measured by fluorescence polarization.

| RNA | Protein | $K_D$ (nM) | Hill slope |
| --- | --- | --- | --- |
| FAM-S1 | SRSF1 | >10000 | - |
| FAM-S2 | SRSF1 | $97.3 \pm 2.5$ | $0.34 \pm 0.01$ |
| FAM-S3 | SRSF1 | $15.9 \pm 1.1$ | $0.37 \pm 0.02$ |
| FAM-H1 | hnRNP A2/B1 | $51.7 \pm 9.0$ | $0.75 \pm 0.09$ |
| FAM-H2 | hnRNP A2/B1 | $134.8 \pm 9.0$ | $0.66 \pm 0.05$ |
| FAM-H3 | hnRNP A2/B1 | $24.0 \pm 8.1$ | $0.52 \pm 0.04$ |

### Analysis of translational repression by flow cytometry

To evaluate whether the repression of TagBFP translation could also be observed as an endpoint assay, we used flow cytometry to analyze different populations. Flow cytometry was previously used to analyze translational repression assays but only by evaluating the fluorescent reporter output.(14) Here, we combined both the sfGFP signal with the TagBFP signal to provide a clear analysis of the fluorescent properties within the bacterial cell population. We used the constructs S4, S5, and S6 in combination with GFP-SRSF1 and compared it to a combination of the same reporters with GFP-PTBP1 to see whether the increase in repression ratio was also reflected in the flow cytometry data. Indeed, with increasing repression ratio we observe a larger difference between the repressed and non-repressed populations (see Fig. 4A-C). In the case of S6 we observed that the populations were almost entirely separated from one another allowing for very clear analysis of the different populations. We also analyzed the population with mutations in the triple repeat sequence (S7-S10) and population changes correlated with the measured repression ratios (see supplemental figures 23 - 26).

**Figure 4:**
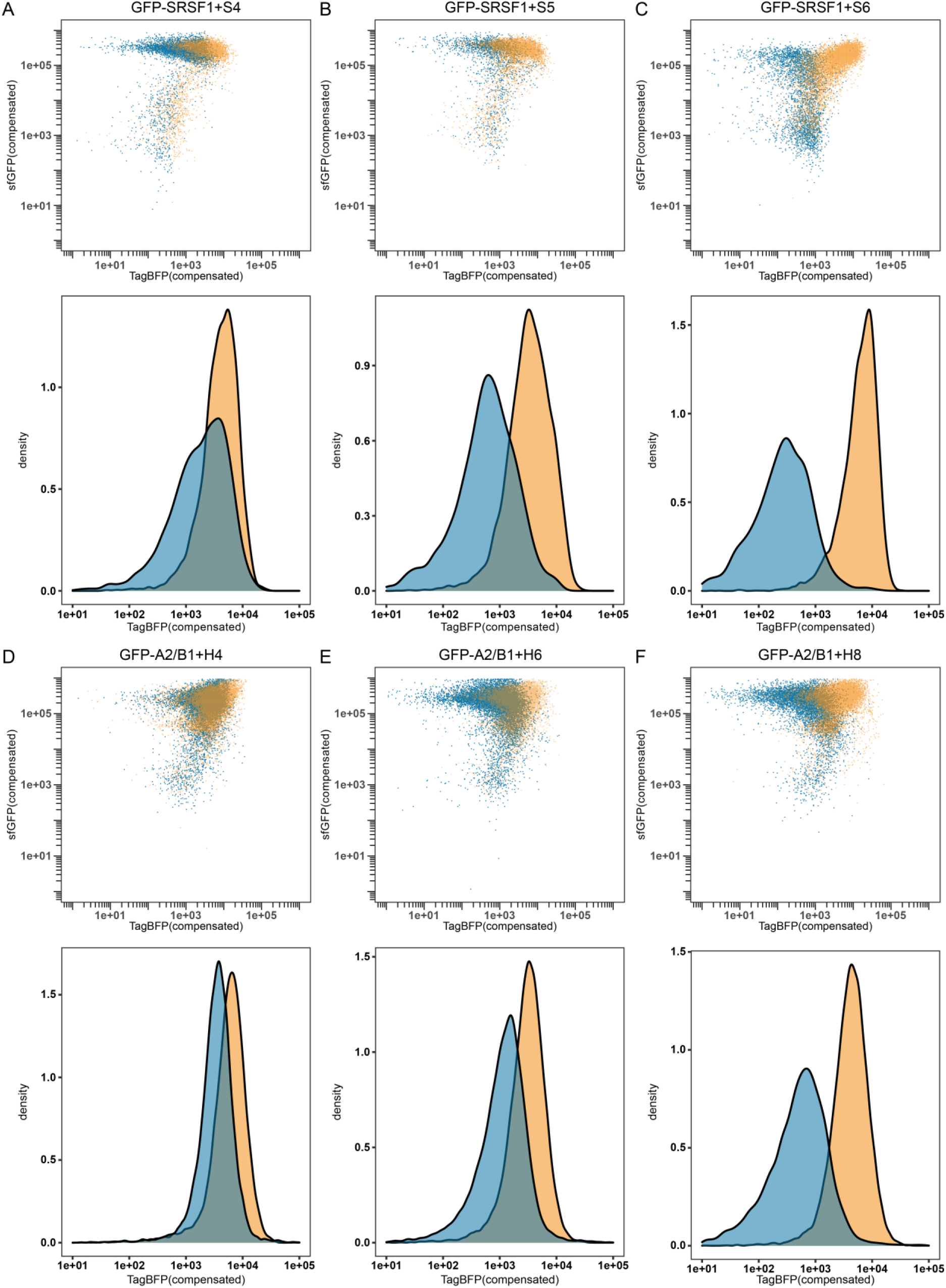
Flow cytometry results for GFP-SRSF1 (blue) or GFP-PTBP1 (orange) in combination with reporter S4 (A), reporter S5 (B) or reporter S6 (C). As well as GFP-A2/B1 (blue) or GFP-PTBP1 (orange) in combination with reporter H4 (D), reporter H6 (E), or reporter H8 (F). Density graphs were produced by analyzing all events >1e05 in the sGFP channel.

We then analyzed several constructs in combination with either GFP-A2/B1 or GFP-PTBP1 and found that similar shifts were observable. Construct H4 showed very little difference between the populations (see Fig. 4D) while the constructs with the higher repression ratios (H6 and H8) showed increasingly larger differences correlating with the repression ratios (see Fig. 4E and F). The results are more obvious when histograms are made of all events of a sufficiently high GFP level (>1*10^5^) to reflect good expression levels of the RBPs (see Fig. 4 and supplemental figures 36-39). These histograms also indicate that a relatively high repression ratio is required for good discrimination between the populations in flow cytometry, further emphasizing that replication of binding sites while avoiding secondary structure formation is essential for a clear read-out. Constructs S7-S10, H5, H7, and H9-H11 were also evaluated and also demonstrated shifts that correlated with the repression ratios (supplemental figures 23-26, 28, 30, and 32-34).

## DISCUSSION

Here we demonstrate that translational repression-based protein-RNA interaction analysis is also feasible using non-structured RNA sequences and can be optimized to analyze protein-RNA pairs with low affinity. The possibility to do so opens this assay up to a wide variety of RNA-binding proteins beyond those that have very high affinity for their target sequences (i.e.: MS2, U1A). When the RNA insert does form a secondary structure in close proximity to the ribosomal binding sites accurate determination of repression values tend to be obstructed due to a reduced basal translation rate. However, by using secondary structure prediction algorithms it is possible to design suitable reporters before cloning and testing in the assay. We demonstrate that repression values determined using time-course assays correlate with end-point analysis using flow cytometry as well as with the affinities measured by fluorescence polarization assays. Simple manipulation of the assay plasmids by straightforward cloning techniques allows the comparative study of variations in RNA sequence or mutations in the RBP. The various constructs described here as well as the design rules provided for the assay allow it to be adapted to other RBPs that bind linear sequences to study the effect of mutation on both the RNA as well as the protein side. The assay is fast and low in cost and avoids the need for the isolation and purification of protein and RNA. It also has the potential to be used in fluorescence assisted cell sorting experiments to select for optimized RNA sequences for a given RBP or vice versa. Furthermore, the effect of reduction in basal reporter protein production rate could potentially be used to investigate the secondary structure of the RNA insert.

## SUPPLEMENTARY DATA

Supplementary Data are available at NAR online.

## ACKNOWLEDGEMENT

We kindly acknowledge the Protein Chemistry Facility of the Max Planck Institute and in particular Jan-Erik Hoffmann for assistance with protein expression and purification. We also thank Pim Huis in ‘t Veld for providing the sfGFP plasmid and Oleg Sitsel for the TagBFP plasmid.

