## Supplemental file 1 for "A translational repression reporter assay for the analysis of RNA-binding protein consensus sites"

### 1. Supplementary tables

Supplemental table 1: Repression ratios SRSF1, hnRNPA2/B1 and PTBP1 in combination with reporter constructs

| Reporter construct | Repression ratio GFP-SRSF1 | Repression ratio GFP-$SRSF1_{mut}$ | Repression ratio GFP-A2/B1 | Repression ratio GFP-PTBP1 |
|---|---|---|---|---|
| S1 (519) | 3.3 ± 0.3 | - | - | 1.5 ± 0.2 |
| S2 (518) | 9.0 ± 2.2 | - | - | 1.7 ± 0.1 |
| S3 (514) | 15.1 ± 2.2 | - | - | 1.7 ± 0.1 |
| S4 (509) | 4.8 ± 1.0 | - | - | 1.4 ± 0.1 |
| S5 (510) | 12.5 ± 1.0 | - | - | 1.5 ± 0.2 |
| S6 (346) | 17.0 ± 0.4 | 1.6 ± 0.2 | - | 1.7 ± 0.1 |
| S7 (511) | 14.2 ± 2.1 | - | - | 1.6 ± 0.2 |
| S8 (512) | 16.1 ± 4.0 | - | - | 1.7 ± 0.2 |
| S9 (513) | 11.8 ± 1.3 | - | - | 1.5 ± 0.3 |
| S10 (542) | 3.2 ± 0.4 | - | - | 1.8 ± 0.3 |
| S0 | 1.2 ± 0.0 | - | - | - |
| H1 (404) | - | - | 5.3 ± 0.4 | 1.6 ± 0.1 |
| H2 (520) | - | - | 2.7 ± 0.4 | 1.3 ± 0.5 |
| H3 (521) | - | - | 5.0 ± 1.1 | 1.3 ± 0.1 |
| H4 (380) | - | - | 3.7 ± 0.2 | 1.3 ± 0.2 |
| H5 (381) | - | - | 2.6 ± 0.6 | 1.3 ± 0.5 |
| H6 (382) | - | - | 4.6 ± 0.7 | 1.4 ± 0.2 |
| H7 (474) | - | - | 4.6 ± 0.2 | 1.1 ± 0.3 |
| H8 (456) | - | - | 6.4 ± 0.1 | 1.4 ± 0.1 |
| H9 (525) | - | - | 5.0 ± 1.7 | 1.3 ± 0.1 |
| H10 (532) | - | - | 4.6 ± 0.3 | 1.2 ± 0.1 |
| H11 (531) | - | - | 6.0 ± 0.5 | 1.1 ± 0.3 |
| H0 | - | - | 1.3 ± 0.2 | - |

Supplemental table 2: List of used plasmids

| Name | Description | Backbone | Insert |
|---|---|---|---|
| S1 | Reporter | pBbA5k[a] | AGAAGAAC |
| S2 | Reporter | pBbA5k | AGAAGAACAGAAGAAC |
| S3 | Reporter | pBbA5k | AGAAGAACAGAAGAACAGAAGAAC |
| S4 | Reporter | pBbA5k | AGAAGAAC |
| S5 | Reporter | pBbA5k | AGAAGAACAGAAGAAC |
| S6 | Reporter | pBbA5k | AGAAGAACAGAAGAACAGAAGAAC |
| S7 | Reporter | pBbA5k | AGAAGUACAGAAGAACAGAAGAAC |
| S8 | Reporter | pBbA5k | AGAAGAACAGAAGUACAGAAGAAC |
| S9 | Reporter | pBbA5k | AGAAGAACAGAAGAACAGAAGUAC |
| S10 | Reporter | pBbA5k | AGAAGUACAGAAGUACAGAAGUAC |
| H1 | Reporter | pBbA5k | AAGGACTAGC |
| H2 | Reporter | pBbA5k | AAGGACTAGCAAGGACTAGC |
| H3 | Reporter | pBbA5k | AAGGACTAGCAAGGACTAGCAAGGACTAGC |
| H4 | Reporter | pBbA5k | AAGGACTAGC |
| H5 | Reporter | pBbA5k | AAGGACTAGCAAGGACTAGC |
| H6 | Reporter | pBbA5k | AAGGACTAGCAAGGACTAGCAAGGACTAGC |
| H7 | Reporter | pBbA5k | AAGGACTAGCGGGAAGGACTAGC |
| H8 | Reporter | pBbA5k | AAGGACTAGCGGGAAGGACTAGC |
| H9 | Reporter | pBbA5k | AAGGACTAGCGGGAAGGACTAGCGGGAAGGACTAGC |
| H10 | Reporter | pBbA5k | AAGCACTAGCGGGAAGGACTAGC |
| H11 | Reporter | pBbA5k | AAGGACTAGCGGGAAGCACTAGC |
| GFP-SRSF1 | RBP | pBbE8c[b] | |
| GFP-hnRNPA2/B1 | RBP | pBbE8c | |
| GFP-SRSF1mut | RBP | pBbE8c | |
| GFP-PTBP1 | RBP | pBbE8c | |

[a]: pBbA5k: Addgene no #35282 and [b]: pBbE8c: Addgene no #35269

Supplemental table 3: List of used RNAs

| Name | Sequence | Modification |
|---|---|---|
| A3 1n | AGAAGAAC | 5'(6-FAM) |
| A3 2n | AGAAGAACAGAAGAAC | 5'(6-FAM) |
| A3 3n | AGAAGAACAGAAGAACAGAAGAAC | 5'(6-FAM) |
| Polypyrimidine | AUUUUUCCAUCUUUGUAUC | 5'(6-FAM) |
| 10 mer 1n | AAGGACUAGC | 5'(6-FAM) |
| 10 mer 2n | AAGGACUAGCAAGGACUAGC | 5'(6-FAM) |
| 10 mer 3n | AAGGACUAGCAAGGACUAGCAAGGACUAGC | 5'(6-FAM) |
| Poly C | CCCCCCCC | 5'(6-FAM) |

Supplemental table 4: List of used proteins

| Name | Protein | Tags | Residues |
|---|---|---|---|
| SRSF1 RRM1+2 | SRSF1 | - | 1-195 |
| MBP-A2B1 | hnRNP A2/B1 | MBP | 1-251 |

### 2. Supplementary figures

#### 2.1 Supplementary figures: Translational repression assay data

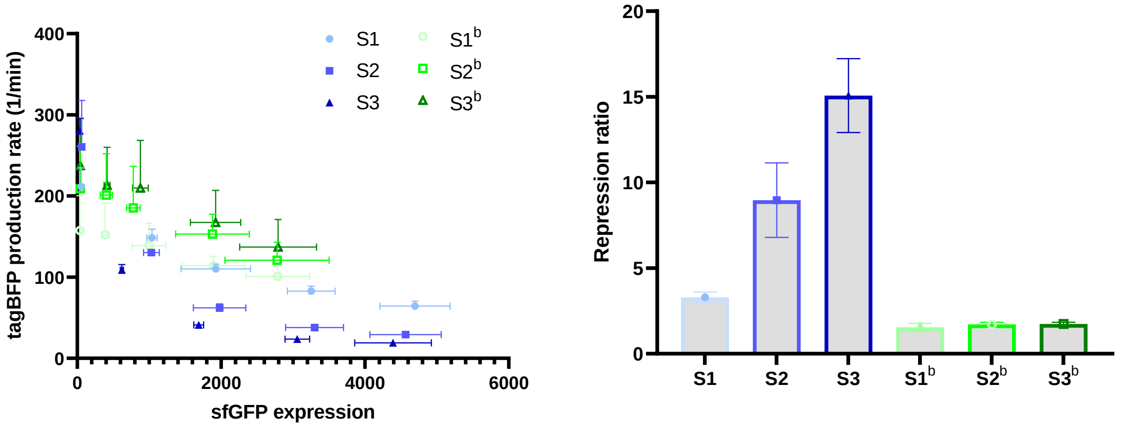

Supplemental figure 1: TRAP assay data of reporter constructs S1-S3. Left: Repression curves for GFP-SRSF1 (blue tones) and GFP-PTBP1 (green tones, indicated by b). Right: Repression ratios for GFP-SRSF1 (blue tones) and GFP-PTBP1 (green tones, indicated by b). Data are mean values (n=2, N=2).

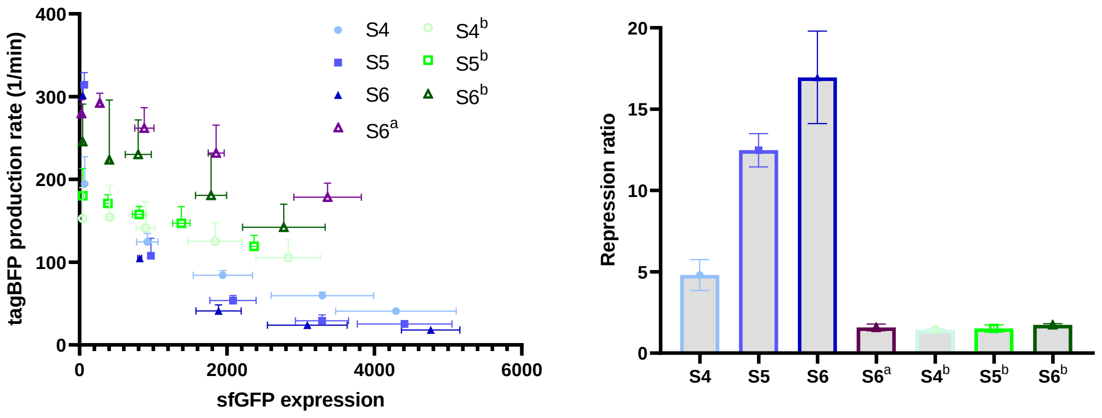

Supplemental figure 2: TRAP assay data of reporter constructs S4-S6. Left: Repression curves for GFP-SRSF1 (blue tones), GFP-SRSF1$_{mut}$ (purple, indicated by a) and GFP-PTBP1 (green tones, indicated by b). Right: Repression ratios for GFP-SRSF1 (blue tones) GFP-SRSF1$_{mut}$ (purple, indicated by a) and GFP-PTBP1 (green tones, indicated by b). Data are mean values (n=2, N=2).

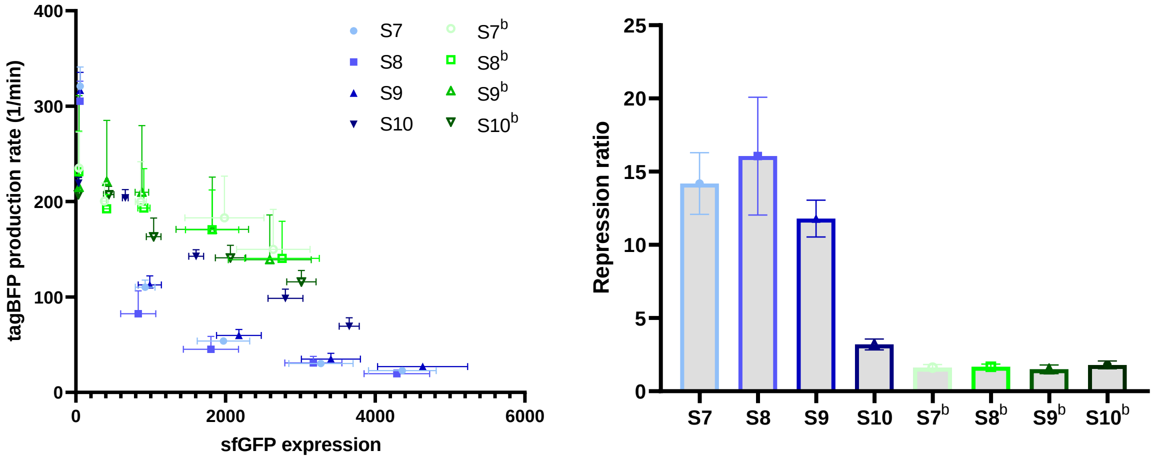

Supplemental figure 3: TRAP assay data of reporter constructs S7-S10. Left: Repression curves for reporters S7-S10 with GFP-SRSF1 (blue tones) and GFP-PTBP1 (green tones). Right: Repression ratios for reporters S7-S10 with GFP-SRSF1 (blue tones) and GFP-PTBP1 (green tones). Data are mean values (n=2, N=2).

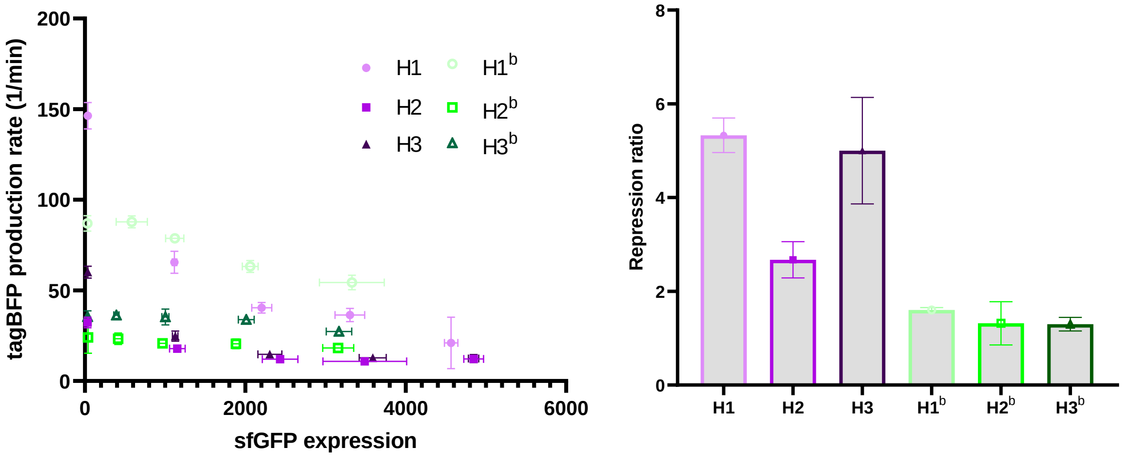

Supplemental figure 4: TRAP assay data of reporter constructs H1-H3. Left: Repression curves for GFP-A2/B1 (pink tones) and GFP-PTBP1 (green tones, indicated by b). Right: Repression ratios for GFP-A2/B1 (pink tones) and GFP-PTBP1 (green tones, indicated by b). Data are mean values (n=2, N=2).

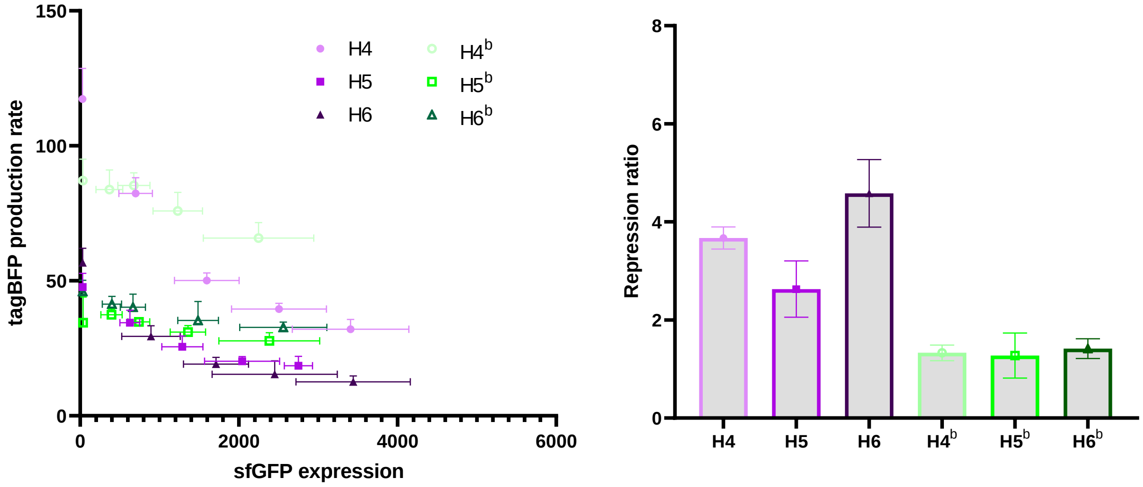

Supplemental figure 5: TRAP assay data of reporter constructs H4-H6. Left: Repression curves for GFP-A2/B1 (pink tones) and GFP-PTBP1 (green tones, indicated by b). Right: Repression ratios for GFP-A2/B1 (pink tones) and GFP-PTBP1 (green tones, indicated by b). Data are mean values (n=2, N=2).

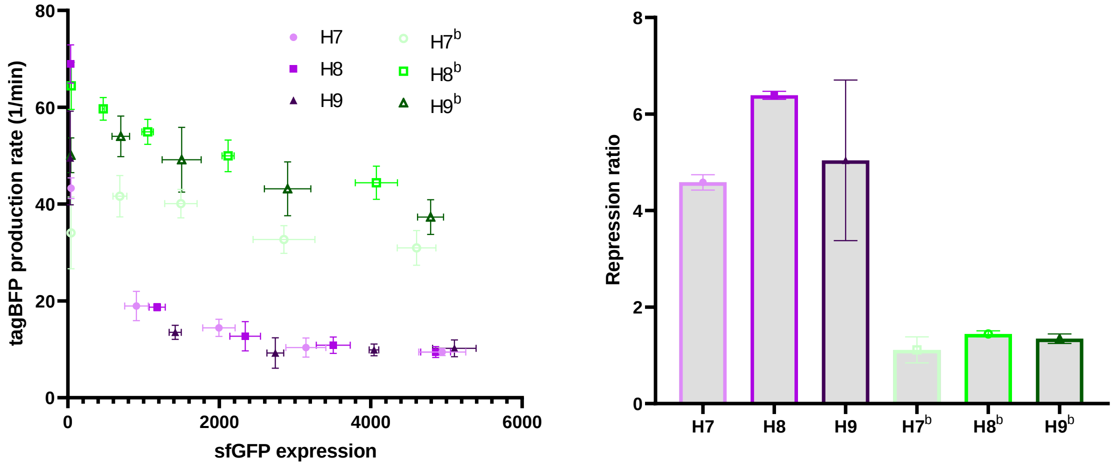

Supplemental figure 6: TRAP assay data of reporter constructs H7-H9. Left: Repression curves for GFP-A2/B1 (pink tones) and GFP-PTBP1 (green tones, indicated by b). Right: Repression ratios for GFP-A2/B1 (pink tones) and GFP-PTBP1 (green tones, indicated by b). Data are mean values (n=2, N=2).

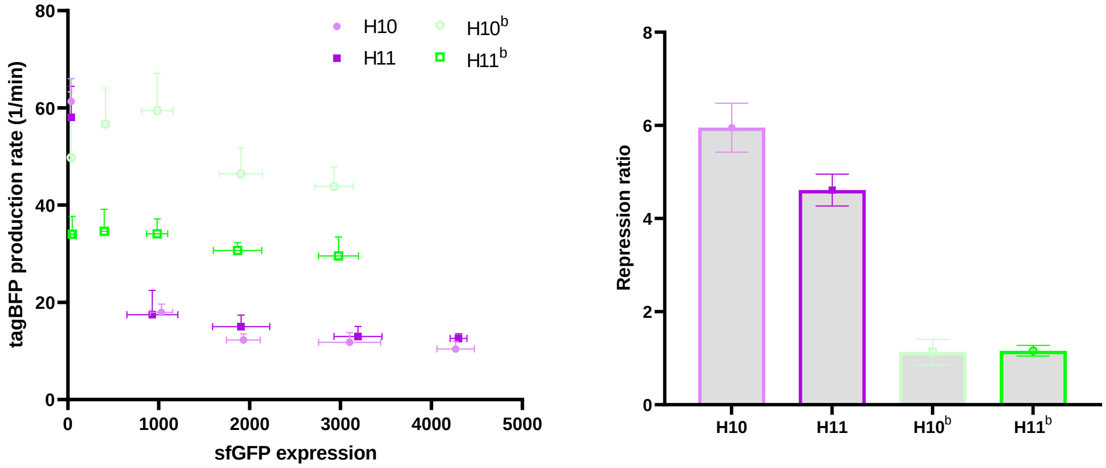

Supplemental figure 7: TRAP assay data of reporter constructs H10 and H11. Left: Repression curves for GFP-A2/B1 (pink tones) and GFP-PTBP1 (green tones, indicated by b). Right: Repression ratios for GFP-A2/B1 (pink tones) and GFP-PTBP1 (green tones, indicated by b). Data are mean values (n=2, N=2).

### 2.2. Supplementary figures: Secondary structure analysis

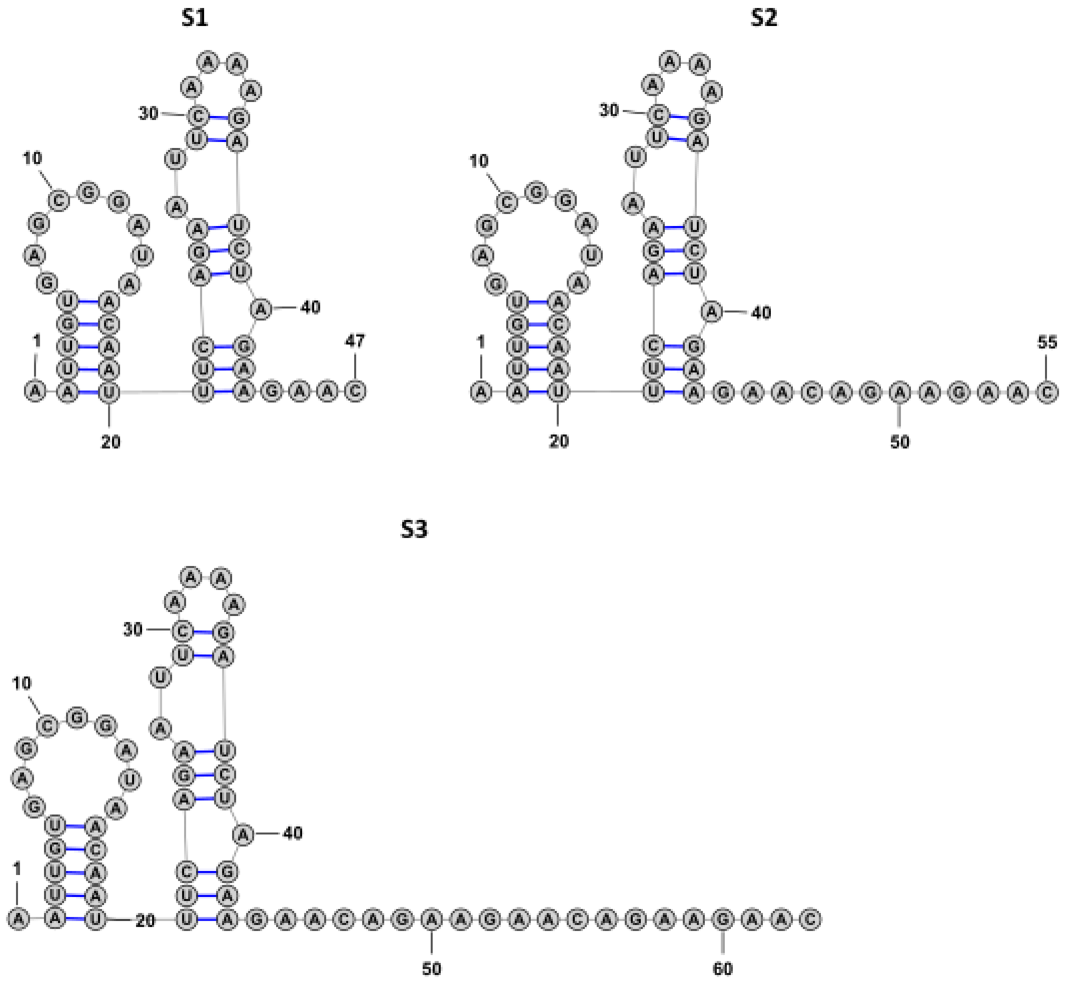

Supplemental figure 8: Secondary structure analysis for reporters S1, S2 and S3 with hairpin structures using RNAfold.

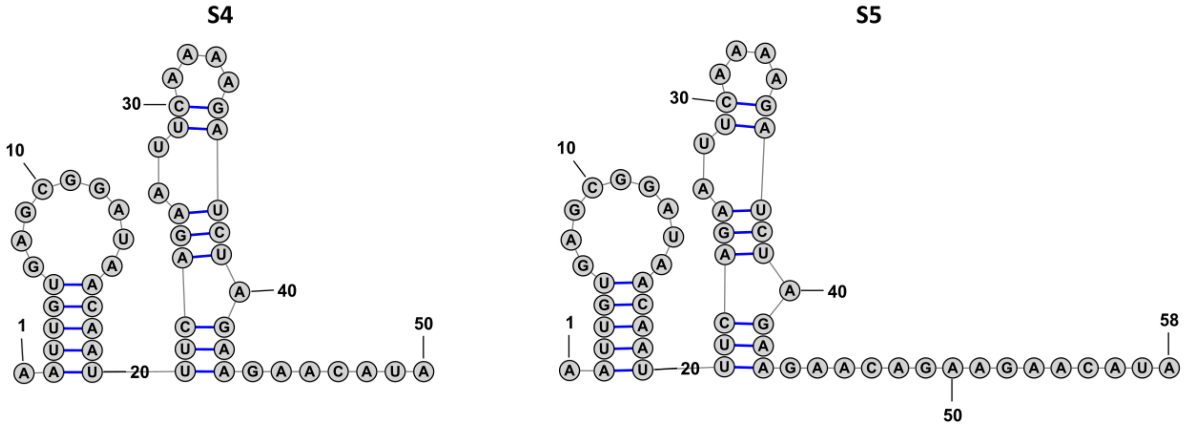

**S6**

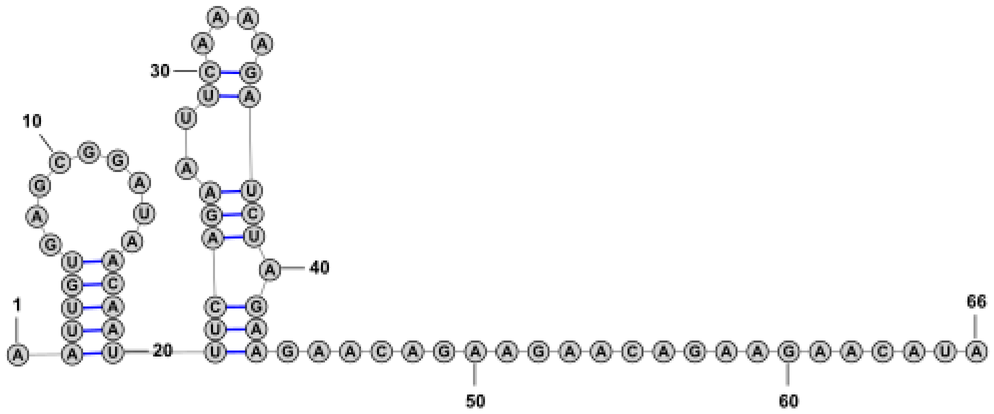

Supplemental figure 9: Secondary structure analysis for reporters S4, S5 and S6 with hairpin structures using RNAfold.

**S7**

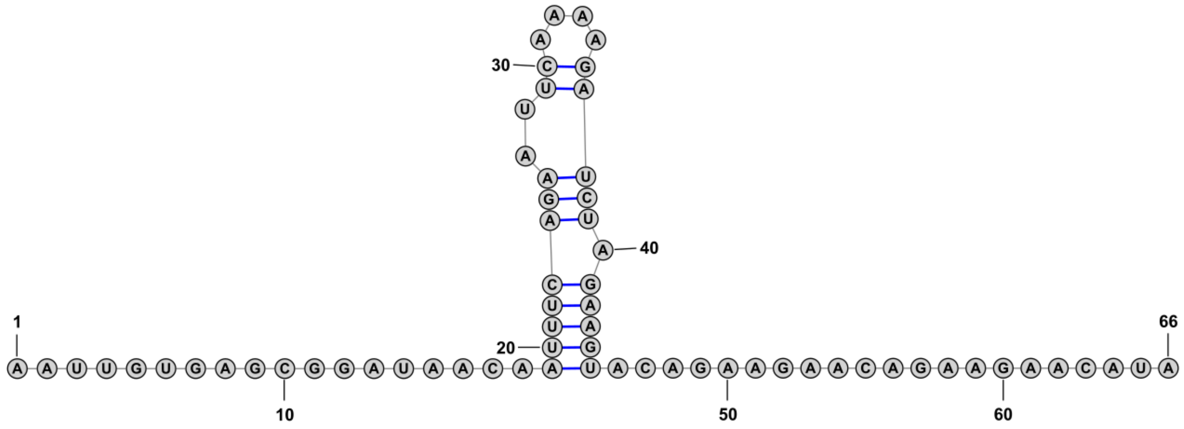

**S8**

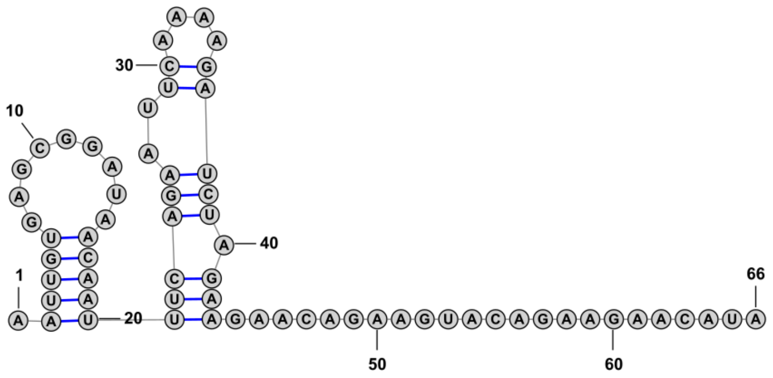

**S9**

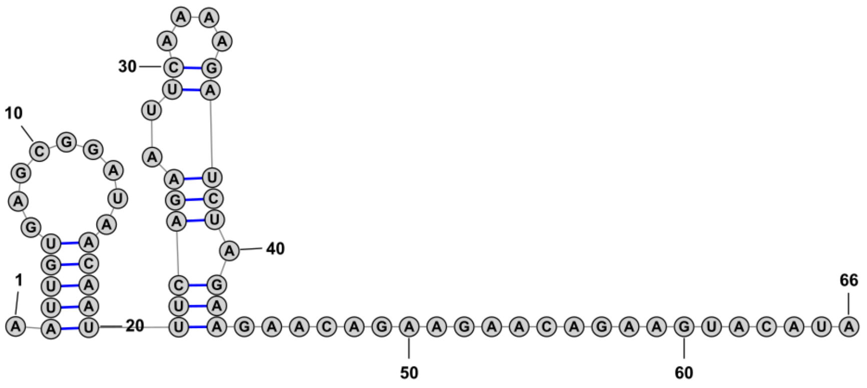

**S10**

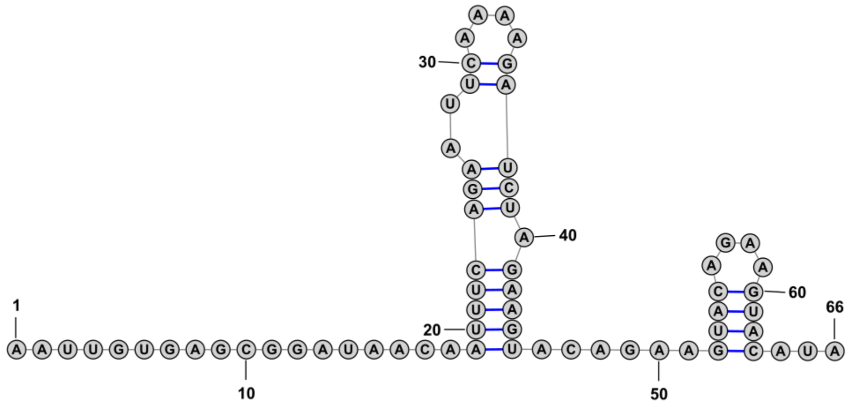

Supplemental figure 10: Secondary structure analysis for reporters S7, S8, S9 and S10 with hairpin structures using RNAfold.

**H1**

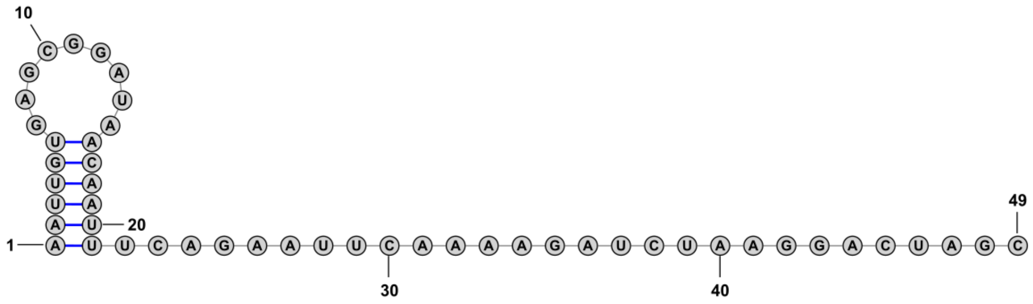

**H2**

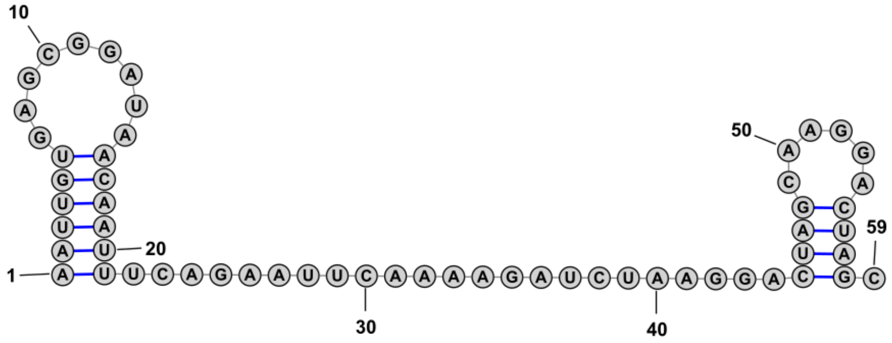

**H3**

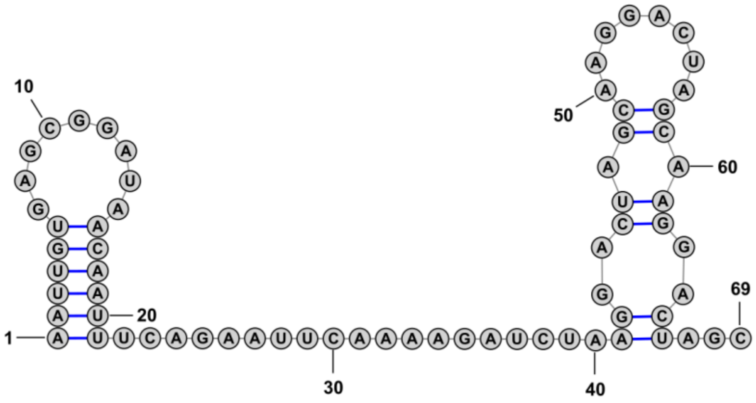

Supplemental figure 11: Secondary structure analysis for reporters H1, H2 and H3 with hairpin structures using RNAfold.

**H4**

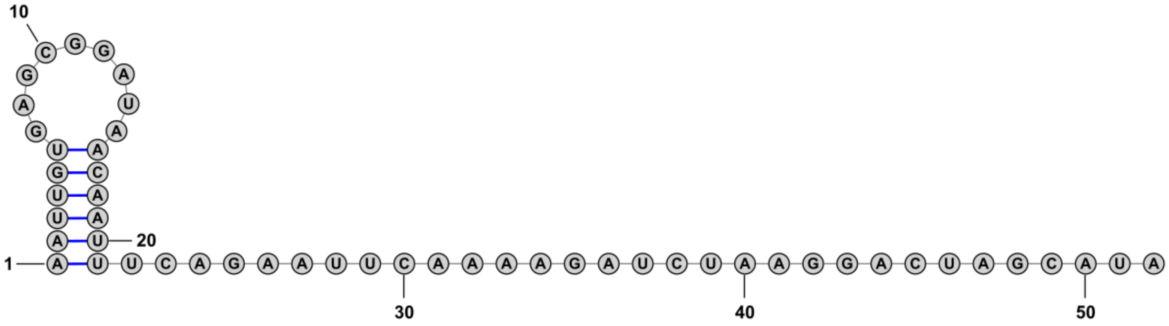

**H5**

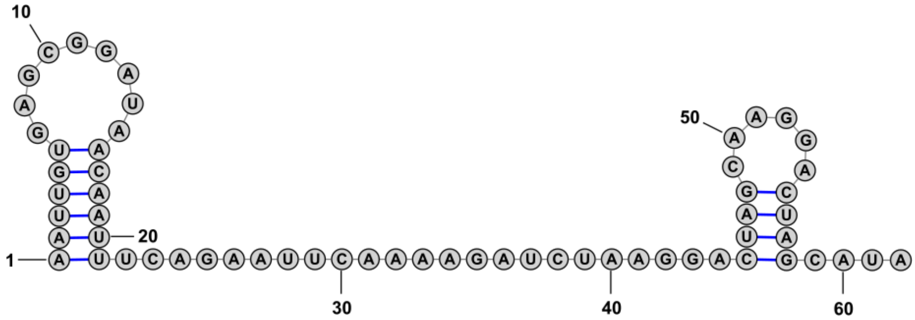

**H6**

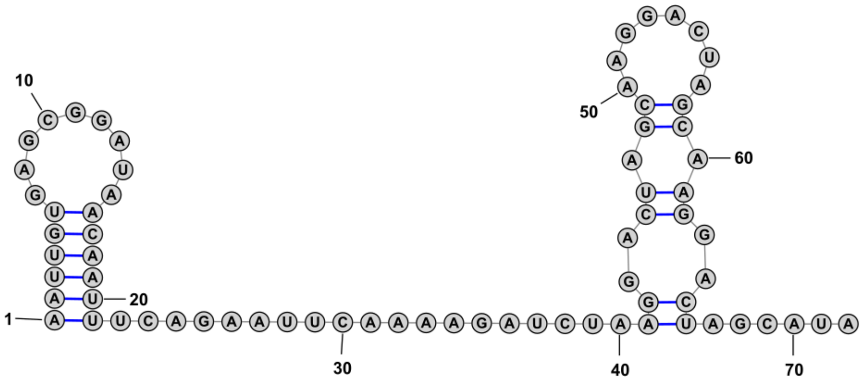

Supplemental figure 12: Secondary structure analysis for reporters H4, H5 and H6 with hairpin structures using RNAfold.

**H7**

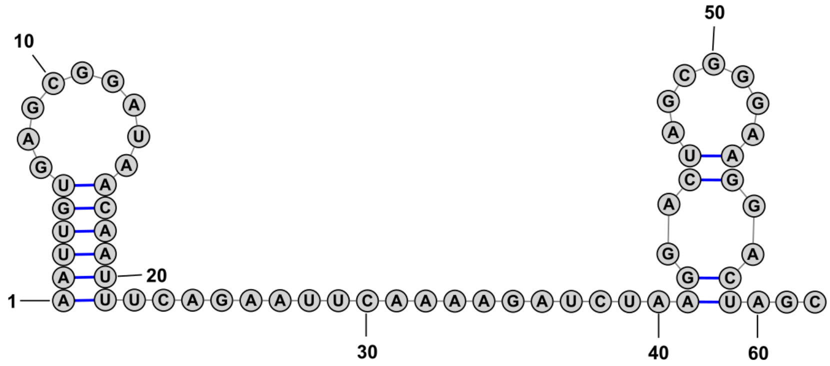

**H8**

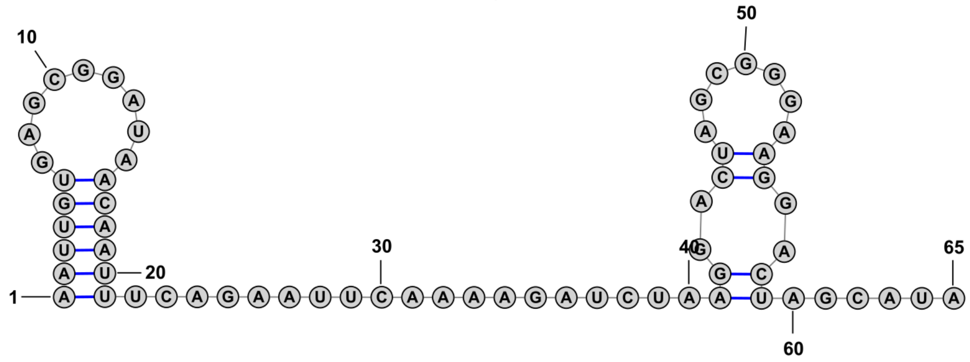

**H9**

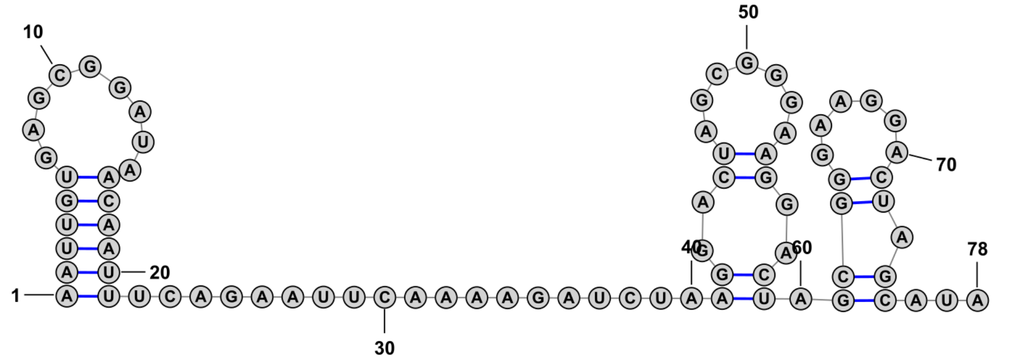

Supplemental figure 13: Secondary structure analysis for reporters H7, H8 and H9 with hairpin structures using RNAfold.

**H10**

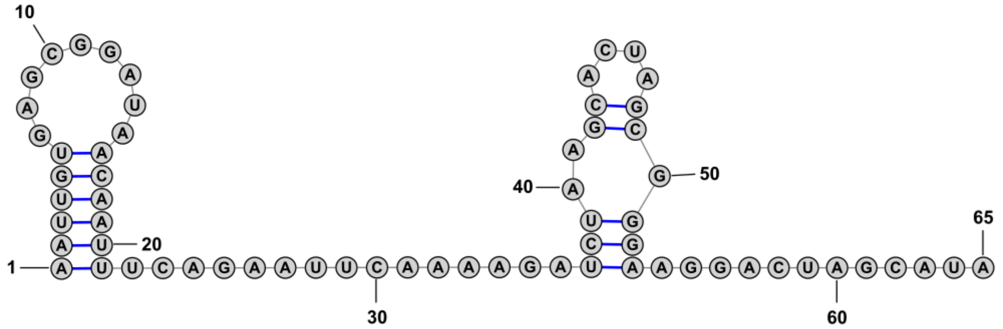

**H11**

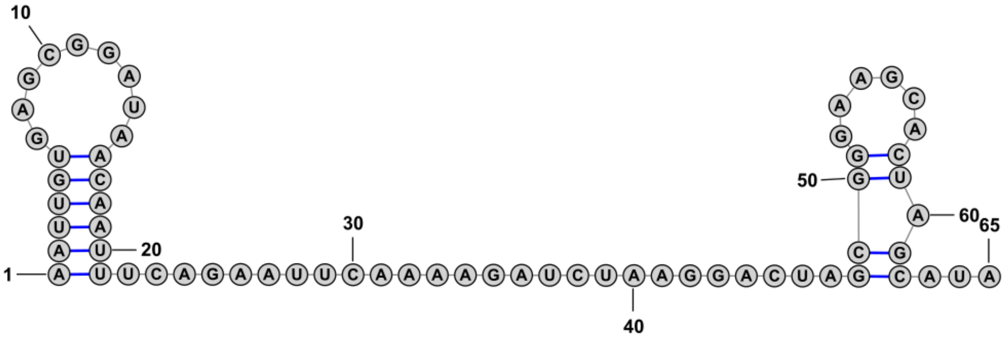

Supplemental figure 14: Secondary structure analysis for reporters H10 and H11 with hairpin structures using RNAfold.

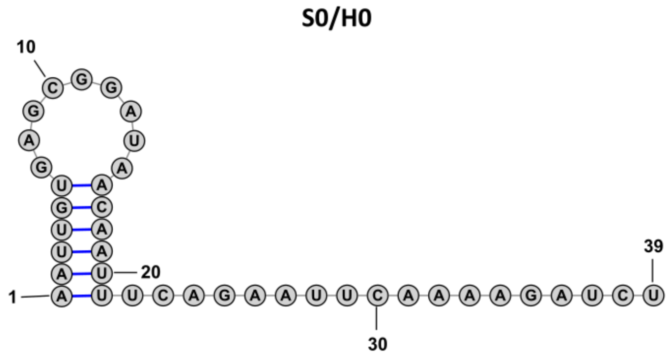

Supplemental figure 15: Secondary structure analysis for reporters S0/H0 with hairpin structures using RNAfold.

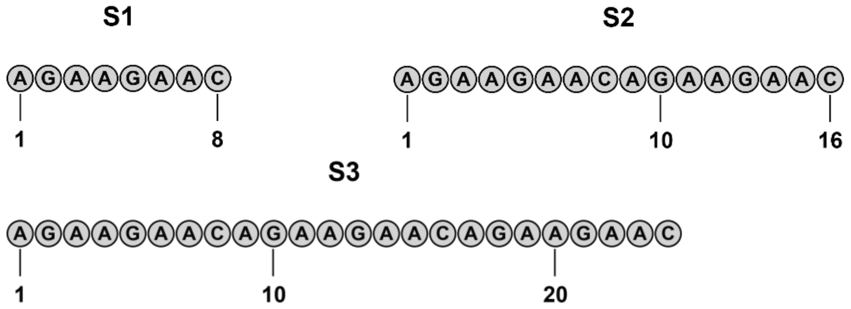

Supplemental figure 16: Secondary structure analysis for RNA sequences FAM-S1 – FAM-S3 used in fluorescence polarization assays using RNAfold.

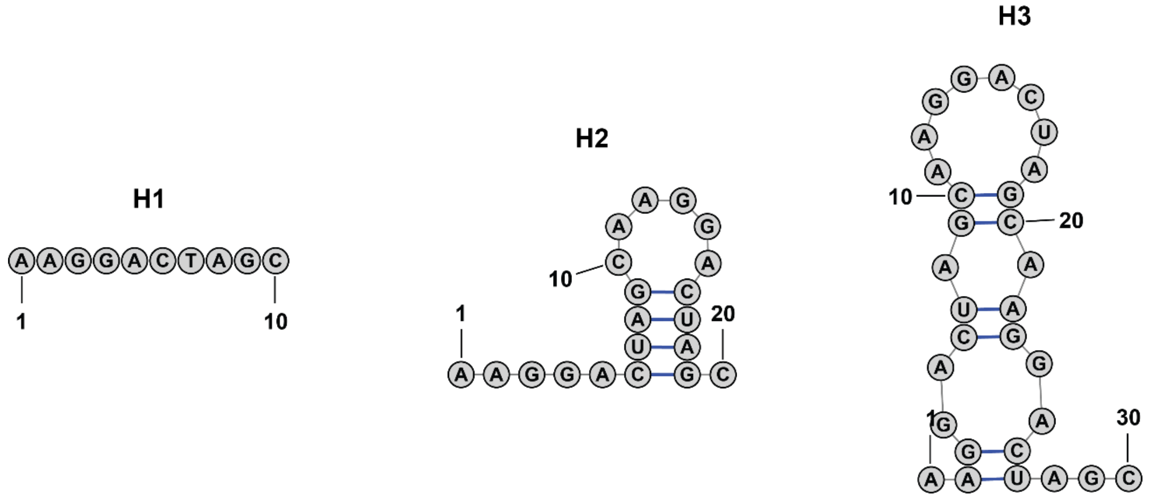

Supplemental figure 17: Secondary structure analysis for RNA sequences FAM-S1 – FAM-S3 used in fluorescence polarization assays using RNAfold.

**2.3. Supplementary figures: Fluorescence polarization**

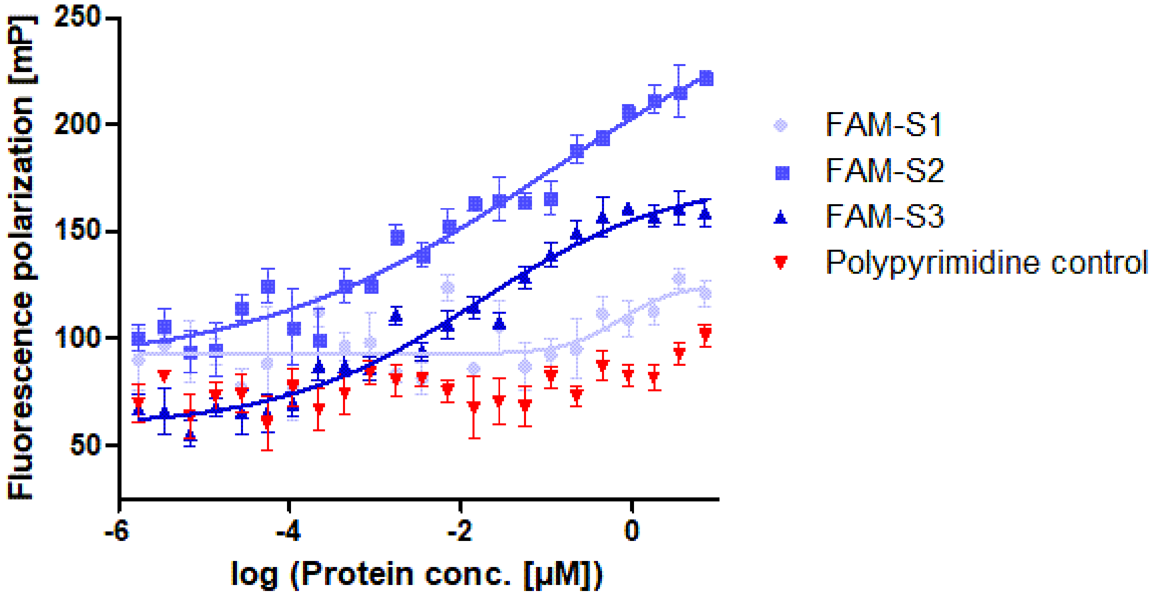

Supplemental figure 18: Fluorescence polarization binding curves of FAM-S1, FAM-S2, FAM-S3, and polypyrimidine RNA with SFSR1.

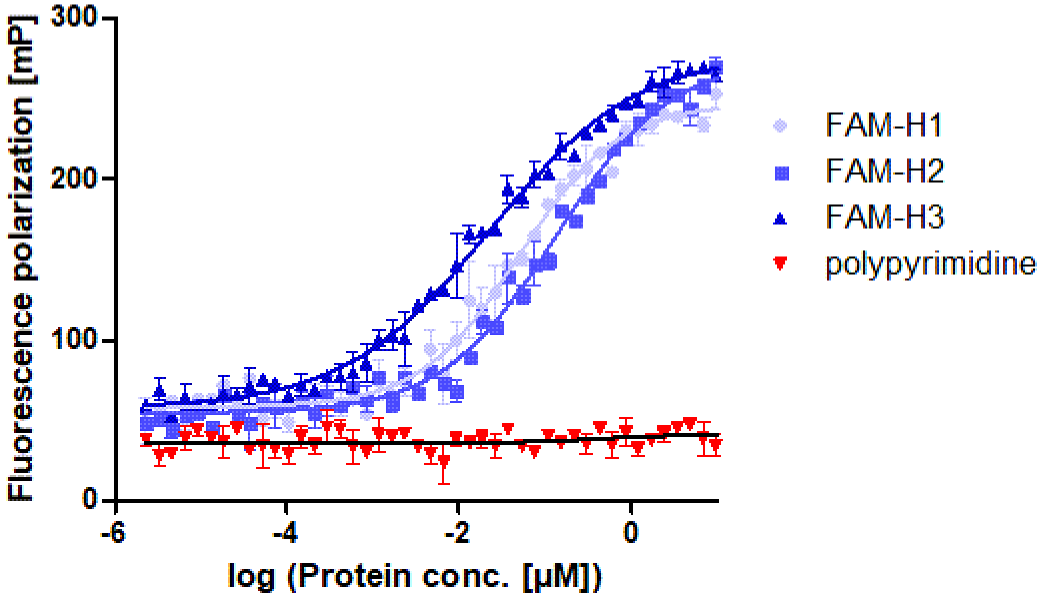

Supplemental figure 19: Fluorescence polarization binding curves of FAM-H1, FAM-H2, FAM-H3, and polypyrimidine RNA with MBP-hnRNP A2/B1.

### 2.4. Supplementary figures: Flow cytometry analysis

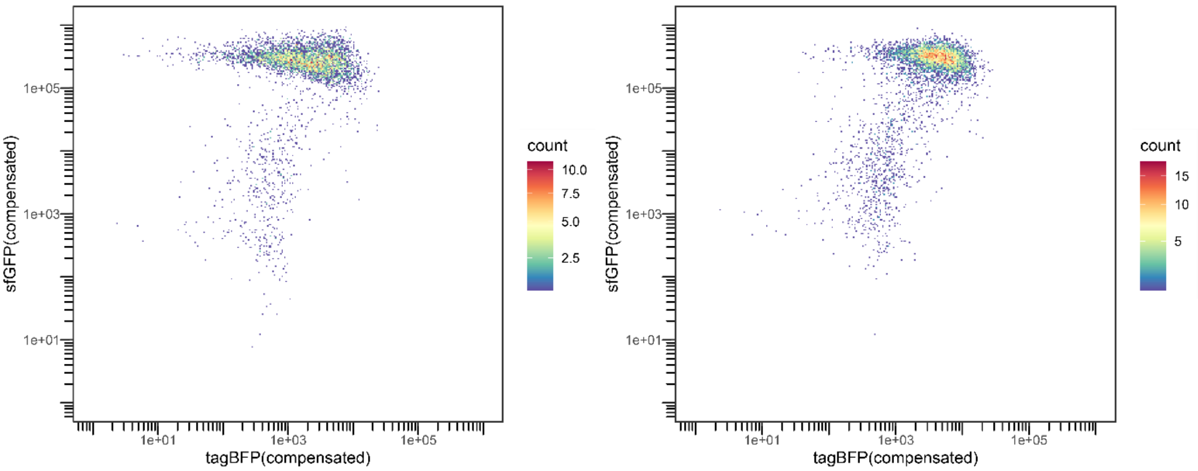

Supplemental figure 20: flow cytometry analysis of reporter S4 with GFP-SRSF1 (left) or GFP-PTBP1 (right)

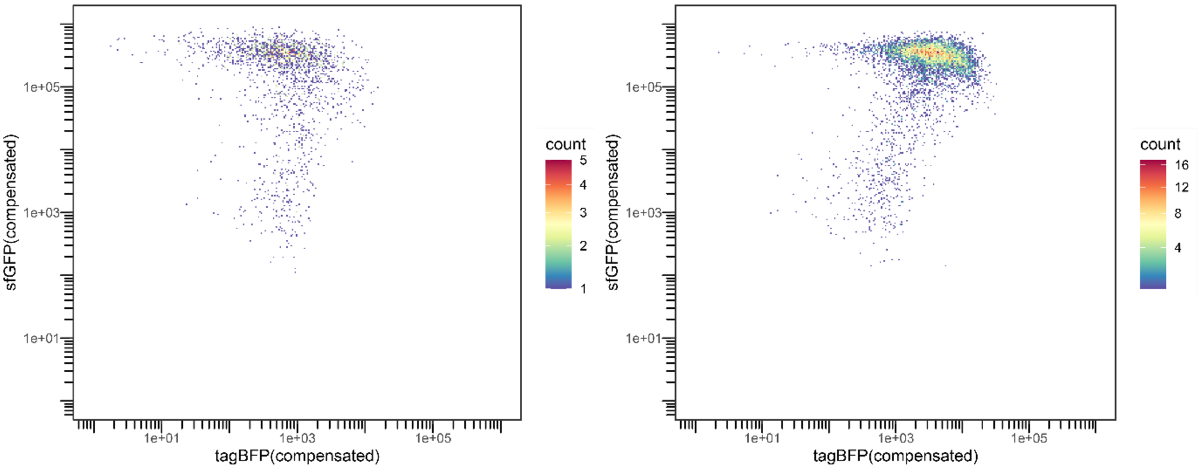

Supplemental figure 21: flow cytometry analysis of reporter S5 with GFP-SRSF1 (left) or GFP-PTBP1 (right)

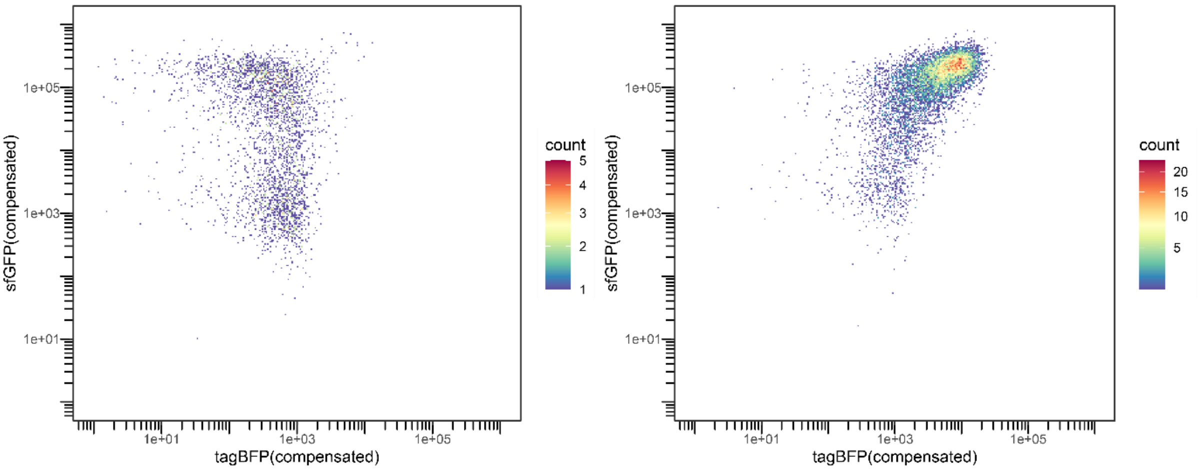

Supplemental figure 22: flow cytometry analysis of reporter S6 with GFP-SRSF1 (left) or GFP-PTBP1 (right)

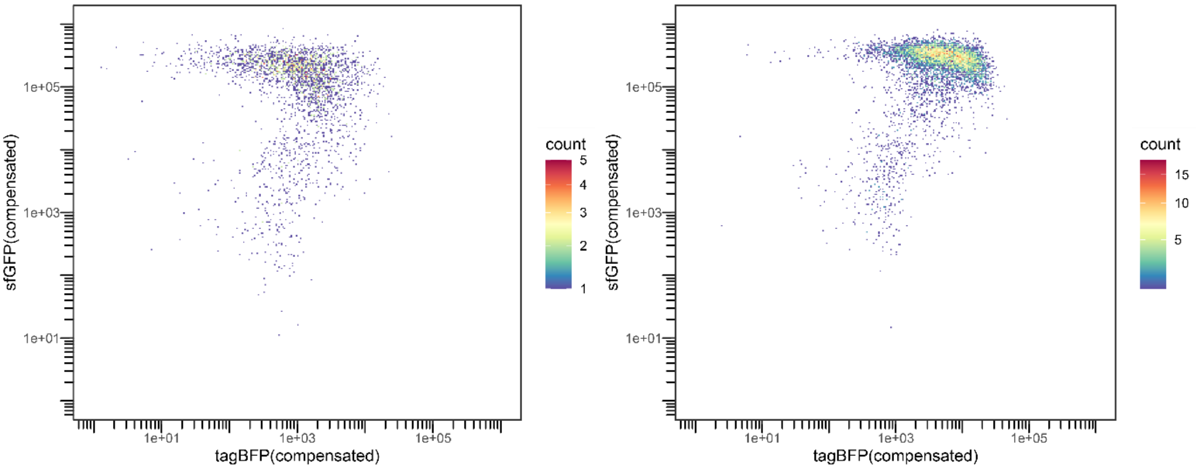

Supplemental figure 23: flow cytometry analysis of reporter S7 with GFP-SRSF1 (left) or GFP-PTBP1 (right)

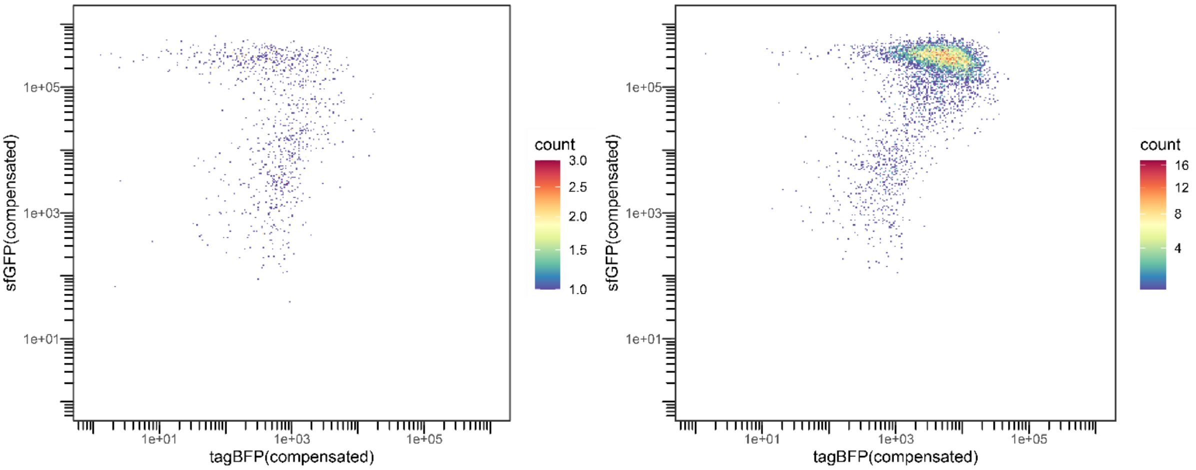

Supplemental figure 24: flow cytometry analysis of reporter S8 with GFP-SRSF1 (left) or GFP-PTBP1 (right)

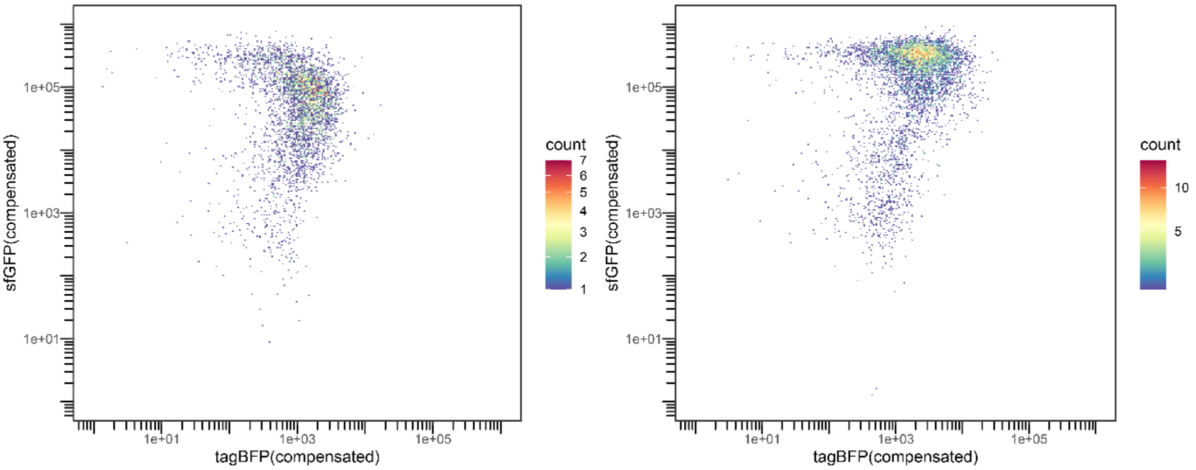

Supplemental figure 25: flow cytometry analysis of reporter S9 with GFP-SRSF1 (left) or GFP-PTBP1 (right)

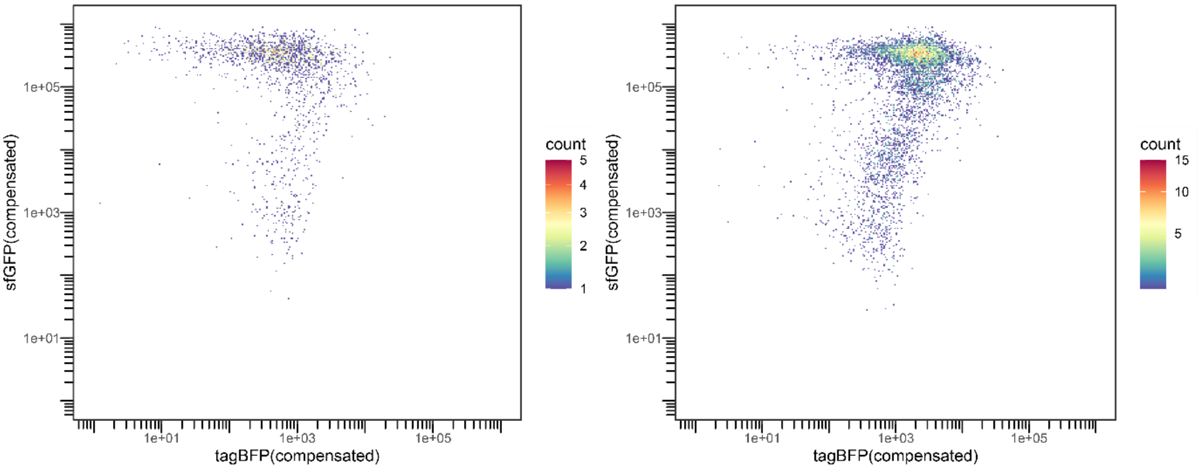

Supplemental figure 26: flow cytometry analysis of reporter S10 with GFP-SRSF1 (left) or GFP-PTBP1 (right)

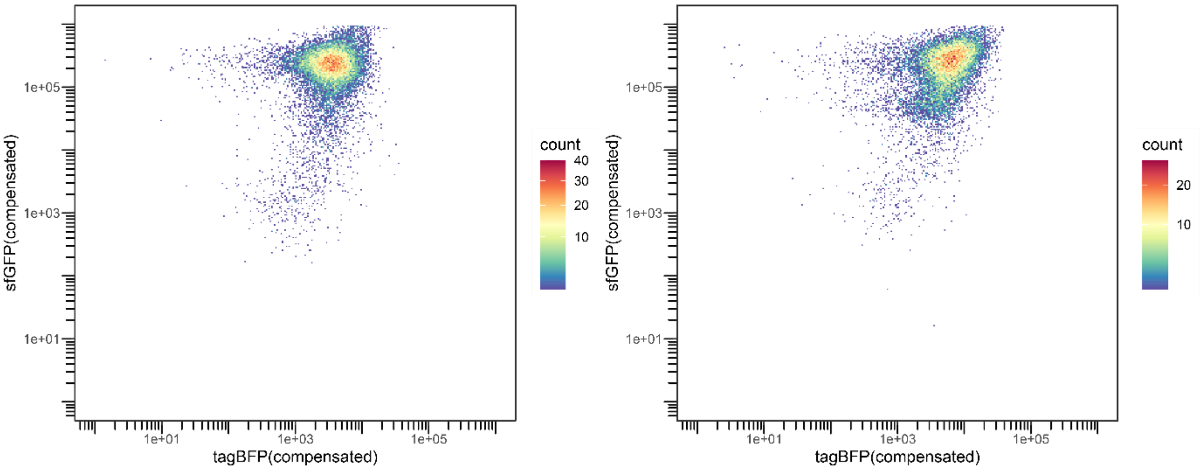

Supplemental figure 27: flow cytometry analysis of reporter H4 with GFP-A2/B1 (left) or GFP-PTBP1 (right)

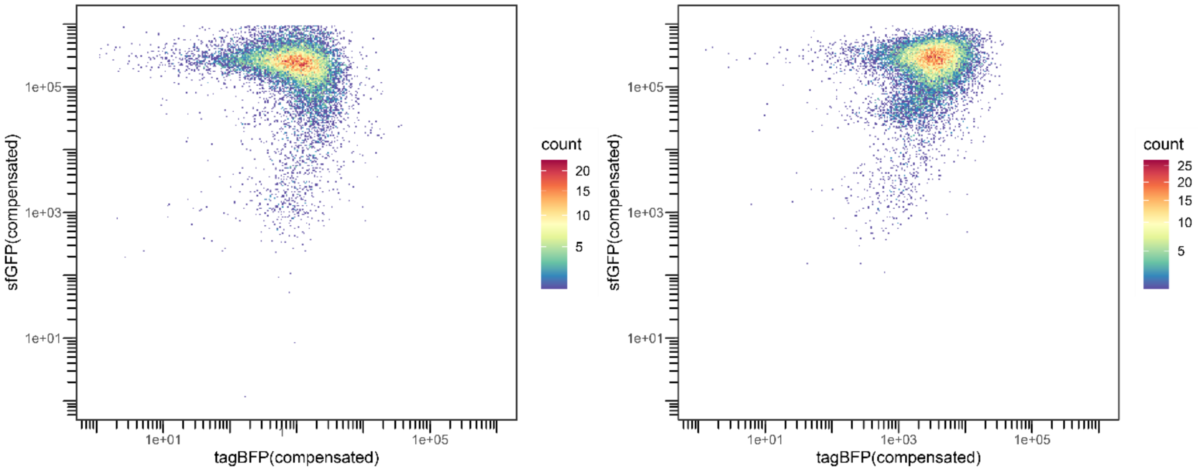

Supplemental figure 28: flow cytometry analysis of reporter H5 with GFP-A2/B1 (left) or GFP-PTBP1 (right)

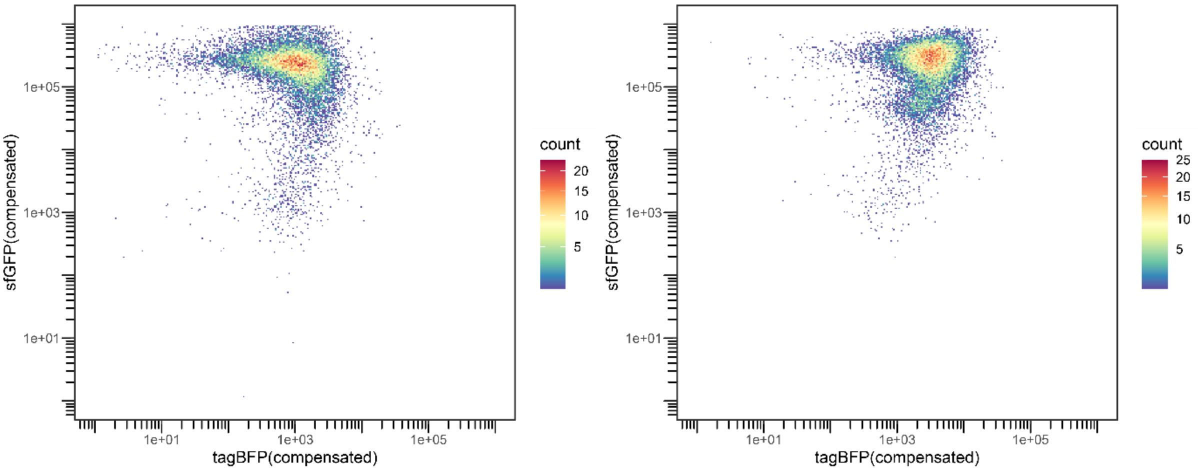

Supplemental figure 29: flow cytometry analysis of reporter H6 with GFP-A2/B1 (left) or GFP-PTBP1 (right)

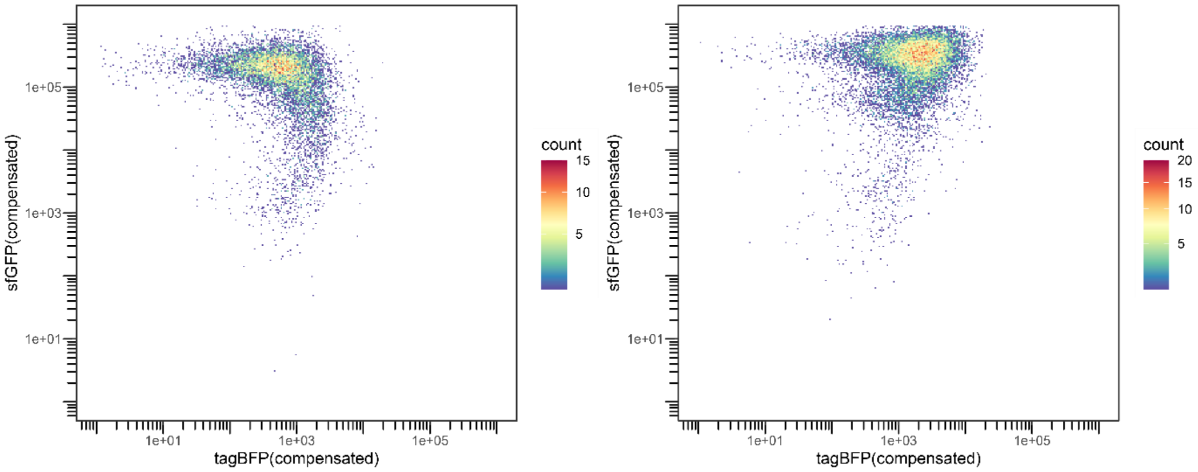

Supplemental figure 30: flow cytometry analysis of reporter H7 with GFP-A2/B1 (left) or GFP-PTBP1 (right)

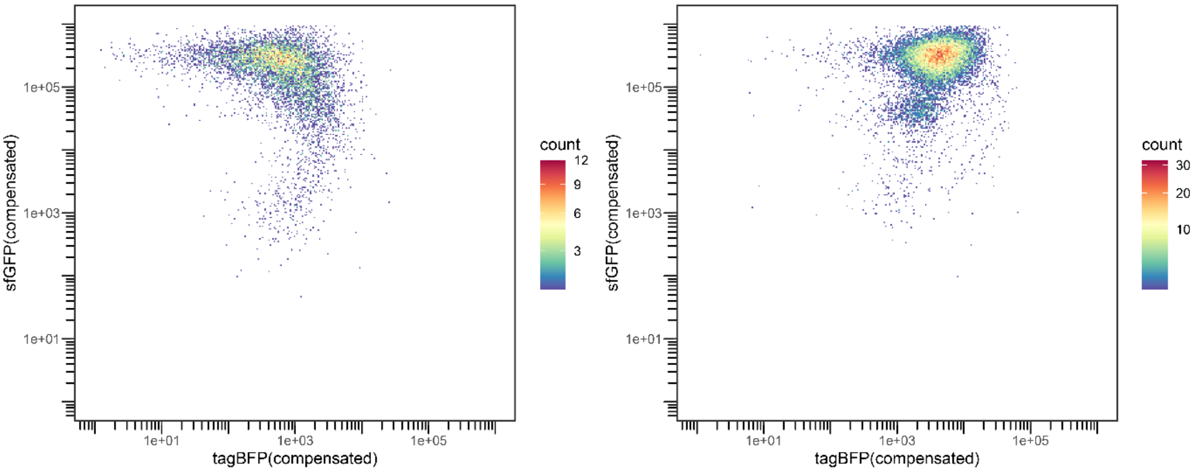

Supplemental figure 31: flow cytometry analysis of reporter H8 with GFP-A2/B1 (left) or GFP-PTBP1 (right)

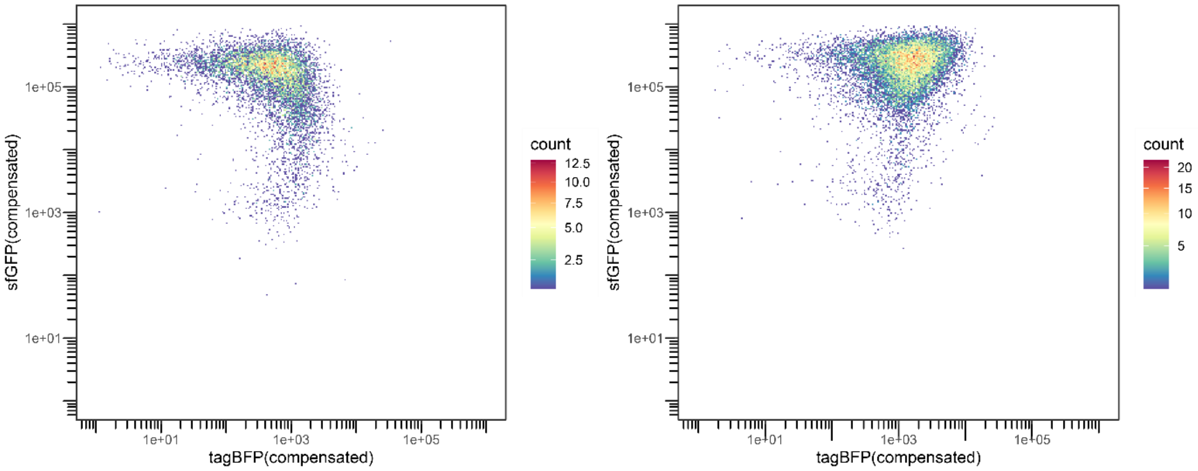

Supplemental figure 32: flow cytometry analysis of reporter H9 with GFP-A2/B1 (left) or GFP-PTBP1 (right)

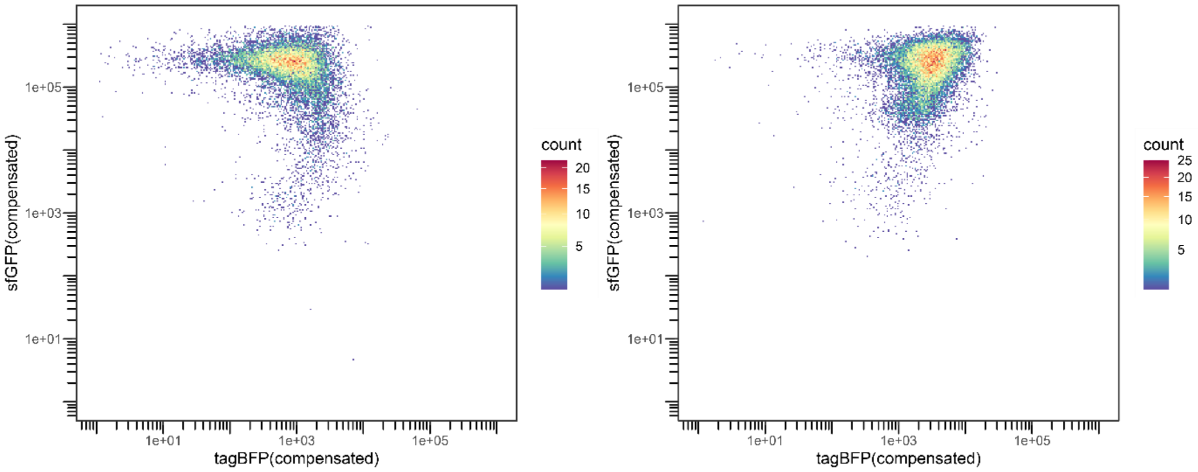

Supplemental figure 33: flow cytometry analysis of reporter H10 with GFP-A2/B1 (left) or GFP-PTBP1 (right)

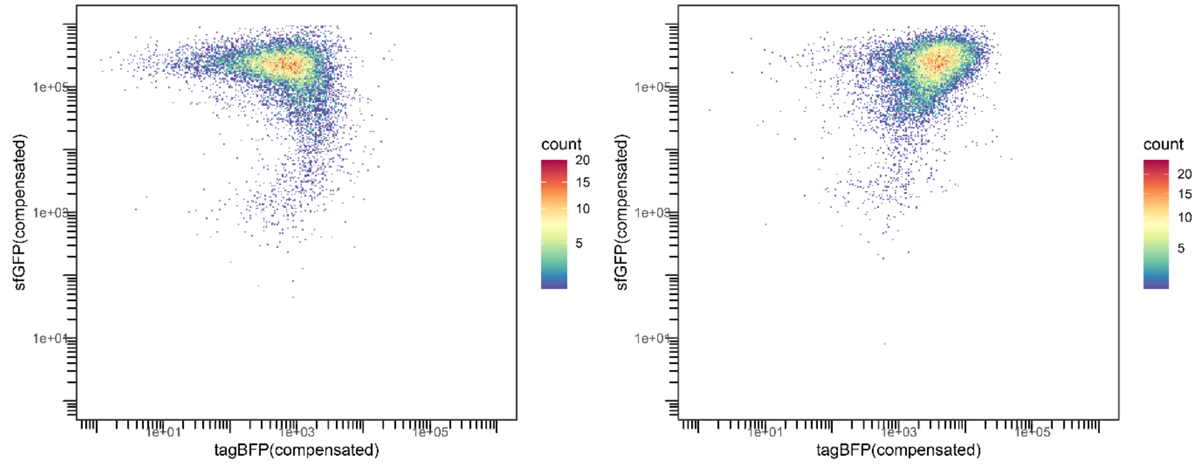

Supplemental figure 34: flow cytometry analysis of reporter H11 with GFP-A2/B1 (left) or GFP-PTBP1 (right)

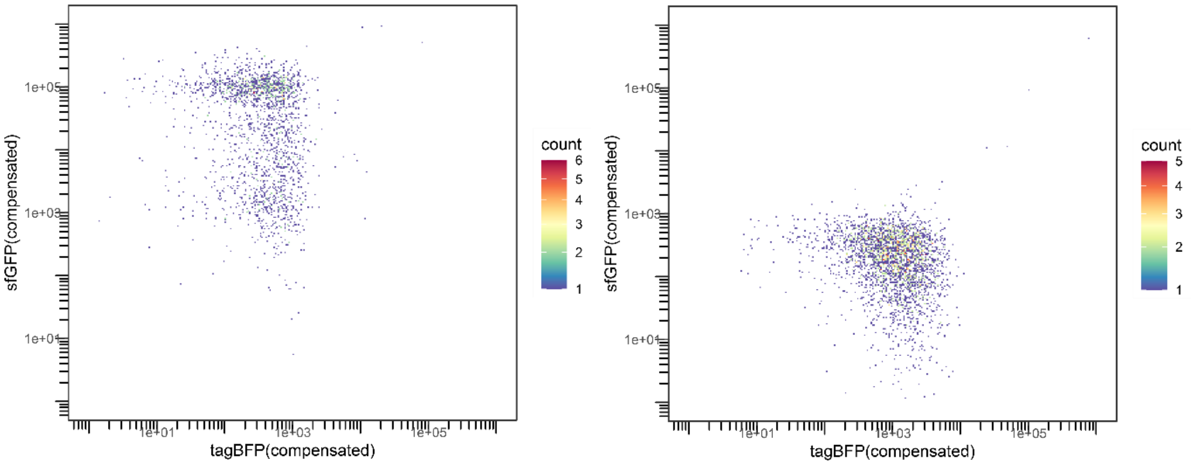

Supplemental figure 35: Flow cytometry results for the single-color controls “GFP only” and “reporter S0/H0” . Top10F’ cells were transformed with either the protein -or the reporter plasmid and grown in medium with the respective antibiotic (chloramphenicol or kanamycin) and induced with either arabinose or IPTG.

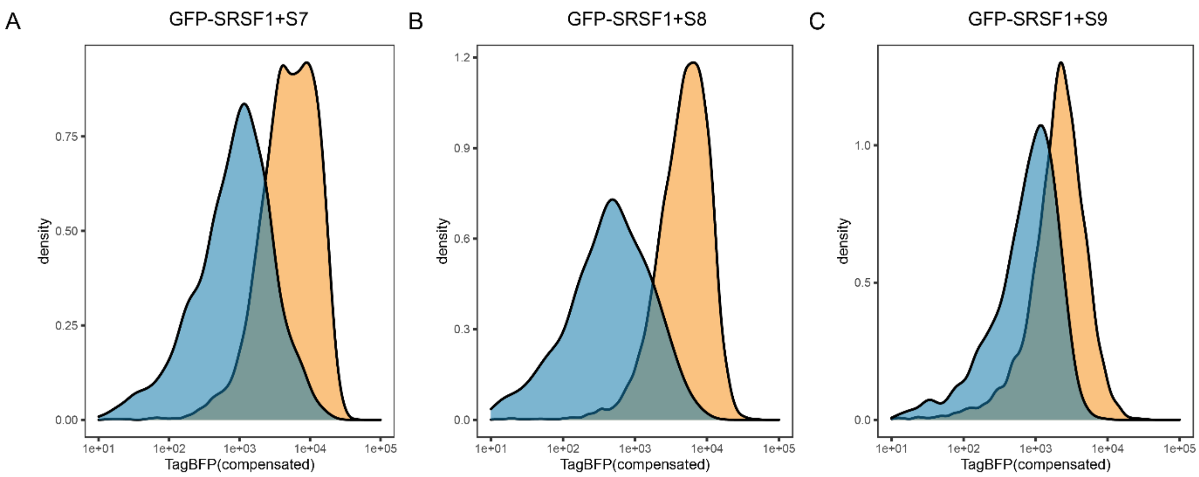

Supplemental figure 36: histograms for reporters S7, S8, and S9 in the presence of GFP-SRSF1 (blue) or GFP-PTBP1 (orange).

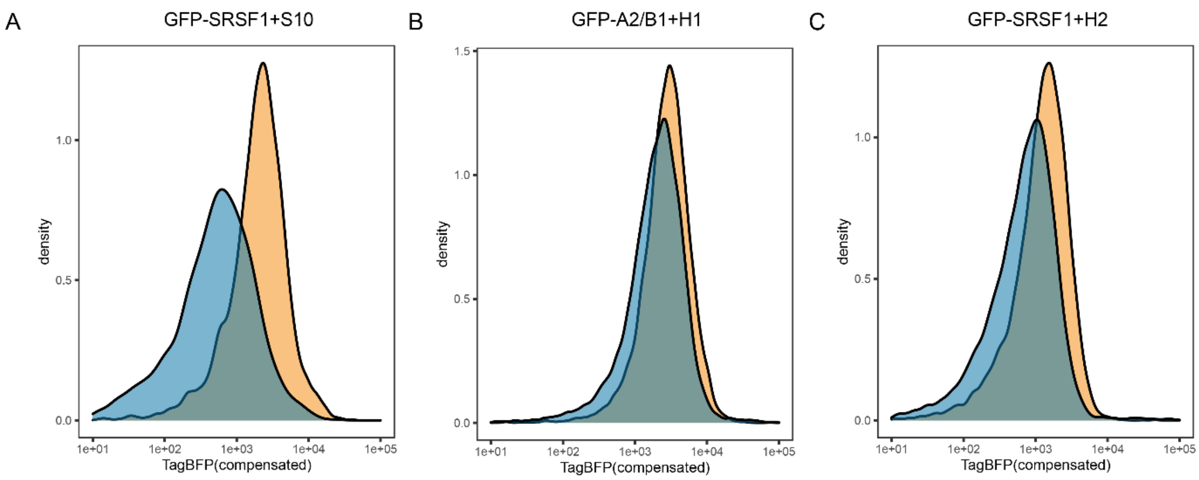

Supplemental figure 37: histograms for reporter S10 in the presence of GFP-SRSF1 (blue) or GFP-

PTBP1 (orange) and reporters H1 and H2 in the presence of GFP-A2/B1 (blue) or GFP-PTBP1 (orange).

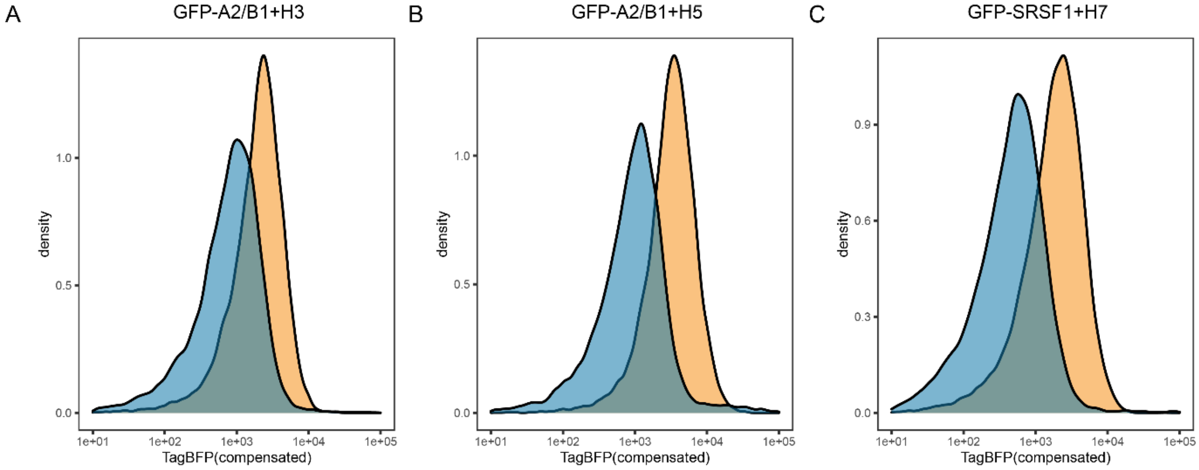

Supplemental figure 38: histograms for reporters H3, H5, and H6 in the presence of GFP-A2/B1 (blue) or GFP-PTBP1 (orange).

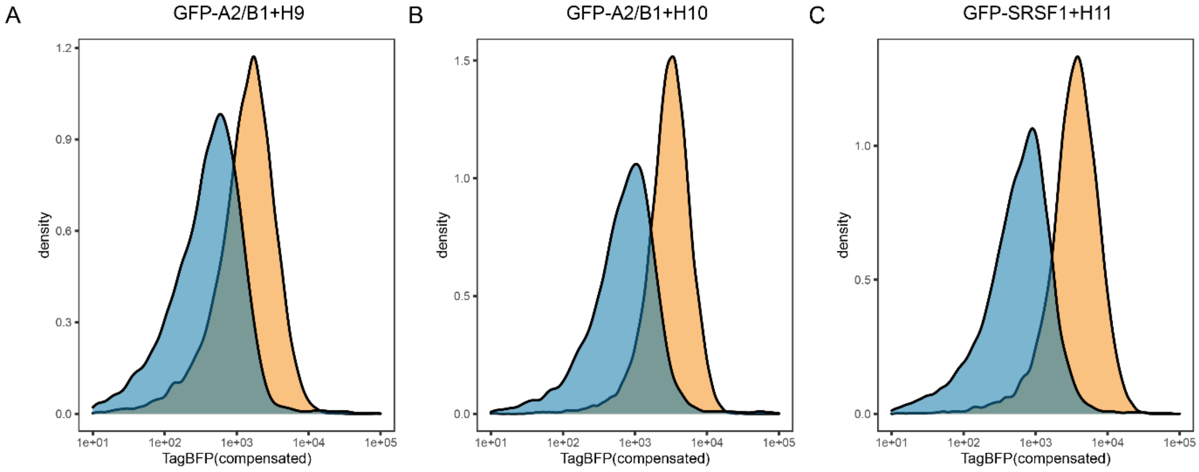

Supplemental figure 39: histograms for reporters H9, H10, and H11 in the presence of GFP-A2/B1 (blue) or GFP-PTBP1 (orange).
